# Small Molecule Activators of B56-PP2A Restore 4E-BP Expression and Function to Suppress Cap-dependent Translation in Cancer Cells

**DOI:** 10.1101/2022.05.24.493272

**Authors:** Michelle A. Lum, Kayla A. Jonas-Breckenridge, Adrian R. Black, Nicholas T. Woods, Caitlin O’Connor, Rita A. Avelar, Analisa DiFeo, Goutham Narla, Jennifer D. Black

## Abstract

Dysregulation of cap-dependent translation is a hallmark of cancer, with key roles in supporting the transformed phenotype. The eIF4E binding proteins (4E-BP1, 2, 3) are major negative regulators of cap-dependent translation that are inactivated in tumors through inhibitory phosphorylation by oncogenic kinases (e.g., mTOR) or by downregulation. Previous studies from our group and others have linked tumor suppressive PP2A family serine/threonine phosphatases to activation of 4E-BP1. Here, we leveraged novel small molecule activators of PP2A (SMAPs) (e.g., DT-061, DT-1154) that are being developed as antitumor agents to (a) explore the role of a subset of B56-PP2As in regulation of 4E-BP activity, and (b) to evaluate the potential of B56-PP2A reactivation for restoring translation control in tumor cells. We show that SMAPs promote PP2A-dependent hypophosphorylation of 4E-BP1/4EBP2 in the presence of active upstream inhibitory kinases (mTOR, ERK, AKT), supporting a role for B56-PP2As as 4E-BP phosphatases. Unexpectedly, DT-061 also led to robust PP2A-dependent upregulation of 4E-BP1, but not 4E-BP2 or 4E-BP3. Cap-binding assays and eIF4E immunoprecipitation showed that SMAP/B56-PP2A blocks the formation of the eIF4F translation initiation complex. Bicistronic reporter assays that directly measure cap-dependent translation activity confirmed the translational consequences of these effects. siRNA knockdown pointed to B56α-PP2A as a mediator of SMAP effects on 4E-BPs, although B56β- and/or B56ε-PP2A may also play a role. 4E-BP1 upregulation involved ATF4-dependent transcription of the 4E-BP1 gene (*EIF4EBP1*) and the effect was partially dependent on TFE3/TFEB transcription factors. Thus, B56-PP2A orchestrates a translation repressive program involving transcriptional induction and hypophosphorylation of 4E-BP1, highlighting the potential of PP2A-based therapeutic strategies for restoration of translation control in cancer cells.

## INTRODUCTION

The activity of the translational machinery is under tight homeostatic control and aberrant activation of translation is seen in a variety of pathologies (1–5). Dysregulated protein synthesis is a central driver of cell transformation and tumor progression (2–5), with key roles in aberrant cell proliferation, survival, angiogenesis, metabolism, immune responses, and chemoresistance (5). Alterations in protein synthesis in tumor cells predominantly occur at the level of translation initiation (1). Hyperactive cap-dependent translation initiation plays a central role in reprogramming the proteome in cancer cells by supporting the translation of a subset of ‘weak’ mRNAs with long and structured 5’ untranslated regions (UTRs). These weak mRNAs typically encode potent oncogenic factors that are required for the transformed phenotype (5–7), including growth factors (e.g., IGF, VEGF), cell cycle regulators (cyclin D, cyclin E), EMT inducers (e.g., Snail), anti-apoptotic proteins (e.g., BCL-xL, MCL1), and transcription factors (e.g., Myc). Oncogenic signaling (Ras/Raf/ERK, PI3K/AKT/mTOR, β-catenin/Myc) hyperactivates cap-dependent translation by disabling members of the eIF4E binding protein (4E-BP) family of translational repressors, which act as a nexus or as “funnel factors” to integrate the effects of signaling pathways on translation (8).

Cap-dependent translation is an exquisitely regulated process that directs ribosomes to the initiation codon in the mRNA template. The initial step in the process is binding of the eukaryotic initiation factor eIF4E to the 5’ m^7^GTP cap of mRNAs (5,9). Cap-bound eIF4E nucleates the assembly of the eIF4F translation initiation complex by engaging the scaffolding protein, eIF4G, and the RNA helicase, eIF4A. eIF4F then recruits the small ribosomal subunit and unwinds secondary structure within the 5’-UTR (5,9). eIF4F assembly is the rate limiting step in cap-dependent translation (5) and is negatively controlled by the 4E-BPs which bind to eIF4E and block its interaction with eIF4G. Three 4E-BPs, 4E-BP1, 4E-BP2, and 4E-BP3, have been identified in mammalian cells, with 4E-BP1 being the most widely expressed and best characterized (1,5).

The activity of 4E-BPs is regulated by hierarchical phosphorylation at conserved sites: Thr37, Thr46, Ser65, and Thr70 (numbering for human 4E-BP1, rodent numbers are lower by one) (10). The hypophosphorylated α and β forms of 4E-BPs interact strongly with eIF4E and block cap-dependent translation, with antiproliferative and tumor suppressive effects (11–13). However, the hyperphosphorylated γ form cannot bind eIF4E and is inactive as a translational repressor. mTOR has been identified as a major kinase mediating inactivation of 4E-BPs in response to growth promoting and oncogenic signaling, phosphorylating all major sites to generate the inactive γ form. However, additional kinases, including ERK, p38, CDK1 and CDK4, also appear to phosphorylate 4E-BPs at specific sites and in certain contexts (8,14,15). Since 4E-BPs act through stoichiometric binding of eIF4E, the relative expression of 4E-BP1 and eIF4E is also critical for control of cap-dependent translation (5,16,17). Thus, in addition to inactivating 4E-BPs by phosphorylation downstream of oncogenic signaling, tumors overcome the repressive function of 4E-BPs by altering the 4E-BP:eIF4E ratio through upregulation of eIF4E and/or downregulation of 4E-BPs (2,3). Loss of 4E-BP function is critical for the transformed phenotype and is also a major mechanism of resistance to clinically relevant targeted agents, including inhibitors of mTOR (18–20), BRAF (21), and MEK (21), which rely on 4E-BP activity for their antitumor effects. Thus, restoration of functional 4E-BP may be a promising therapeutic strategy with potential for overcoming drug resistance. However, relatively little is known regarding the phosphatases that activate the 4E-BPs or the mechanisms that regulate the expression of these critical translation repressors.

Studies by our group and others support a role for the serine/threonine phosphatase, protein phosphatase 2A (PP2A), as a 4E-BP1 phosphatase. We have determined that the anti-proliferative effects of PKCα signaling involve PP2A-mediated dephosphorylation and activation of the translation inhibitory functions of 4E-BP1 (22). Similarly, FGF-induced growth arrest in chondrocytes has been linked to PP2A-mediated activation of 4E-BP1 and inhibition of protein synthesis (23). The ability of PP2A to regulate 4E-BPs is further supported by studies showing that the catalytic subunit of PP2A (PP2A-C) can dephosphorylate 4E-BP1 *in vitro* (24). PP2A is a family of heterotrimeric Ser/Thr phosphatases consisting of a scaffolding A subunit (Aα, Aβ), a catalytic C subunit (Cα, Cβ), and one of 16 regulatory B subunits (25–27). B subunits, which are categorized into four structurally distinct classes, B55 (B), B56 (B’), PR72 (B’’), and Striatins (B’’’), are the major determinants of substrate selectivity, and their diversity allows PP2A to dephosphorylate a broad spectrum of proteins. Thus, PP2A comprises a family of over 60 phosphatases that differ in their substrate specificity and regulation (28). PP2A activity is widely considered a tumor suppressor (28–30), with B56 (α, β, γ, δ, ε) containing holoenzymes emerging as key players in tumor suppression (31,32), and inactivation of PP2A is essential for malignant transformation (31,33,34). Tumors inactivate PP2A by multiple mechanisms, including expression of endogenous inhibitors (e.g., SET, CIP2A), changes in C-subunit post-translational modification (e.g., L309 carboxymethylation), somatic mutation, and decreased expression of individual subunits (31,33–35). While increasing evidence points to PP2A as a regulator of 4E-BP function, the precise heterotrimers involved and the mechanisms underlying PP2A-mediated 4E-BP control remain to be determined.

In this study, we leveraged recently developed small molecule activators of PP2A (SMAPs) (29,36–39) to explore the role of B56-containing PP2As in regulation of 4E-BP activity and to evaluate the potential of PP2A reactivation for restoring translation control in tumor cells. SMAPs activate PP2A by acting as a molecular “glue” to enhance the accumulation of selected PP2A heterotrimers. Extensive structural and functional studies have determined that SMAPs selectively stabilize PP2A heterotrimers through a binding pocket that can accommodate B56α, B56β and B56ε, but not other B56 proteins (B56γ or B56δ) or members of other B subunit families (37). Using SMAPs, we identify B56-containing PP2A enzymes as activators of 4E-BP1 and 4E-BP2 in multiple cell types. Our studies also led to the unexpected finding that B56-PP2A activation promotes transcriptional upregulation of 4E-BP1 via ATF4 and TFE3/TFEB. Together, our findings support the ability of B56-PP2A to oppose all mechanisms used by tumor cells to disable 4E-BP, thus highlighting the potential of PP2A based therapeutic strategies for restoration of translation control in tumors.

## RESULTS

### SMAPs hypophosphorylate and upregulate 4E-BP1

Tumor cells disable 4E-BP1 function through several mechanisms (2,5,15,17,40–42). As shown by analysis of samples from The Cancer Genome Atlas (TCGA) and the Clinical Proteomic Tumor Analysis Consortium (CPTAC), many tumors overexpress 4E-BP1 mRNA (Fig. 1a) and protein (Fig. 1b) (see arrowheads in Fig. 1a,b for examples); however, loss of 4E-BP1 function is achieved in these tumors by inhibitory phosphorylation downstream of oncogenic kinases such as mTOR and ERK (43–46). Thus, expression of high levels of inactive, hyperphosphorylated 4E-BP1 is a characteristic of many cancers. Tumors can also escape translational control by suppressing the expression of 4E-BP1 mRNA (Fig. 1a, arrows) and protein (Fig. 1b, arrows). 4E-BP1 deficiency is a characteristic of pancreatic tumors (PDACs) (Fig. 1a,b, arrows) (2), as confirmed by our proteomic analysis of primary and metastatic PDACs (Fig. 1c) and by Western blot analysis of PDAC cell lines (Fig. 1d). A survey of cancer cell lines further revealed that deficiency of 4E-BP1 protein is commonly seen in colorectal cancer (CRC) (Fig. 1d). In contrast, consistent with TCGA and CPTAC data (Fig.1a,b), the two endometrial cancer (EC) cell lines analyzed (SNG-M and Ishikawa) expressed relatively high levels of 4E-BP1 protein.

**Figure 1.**
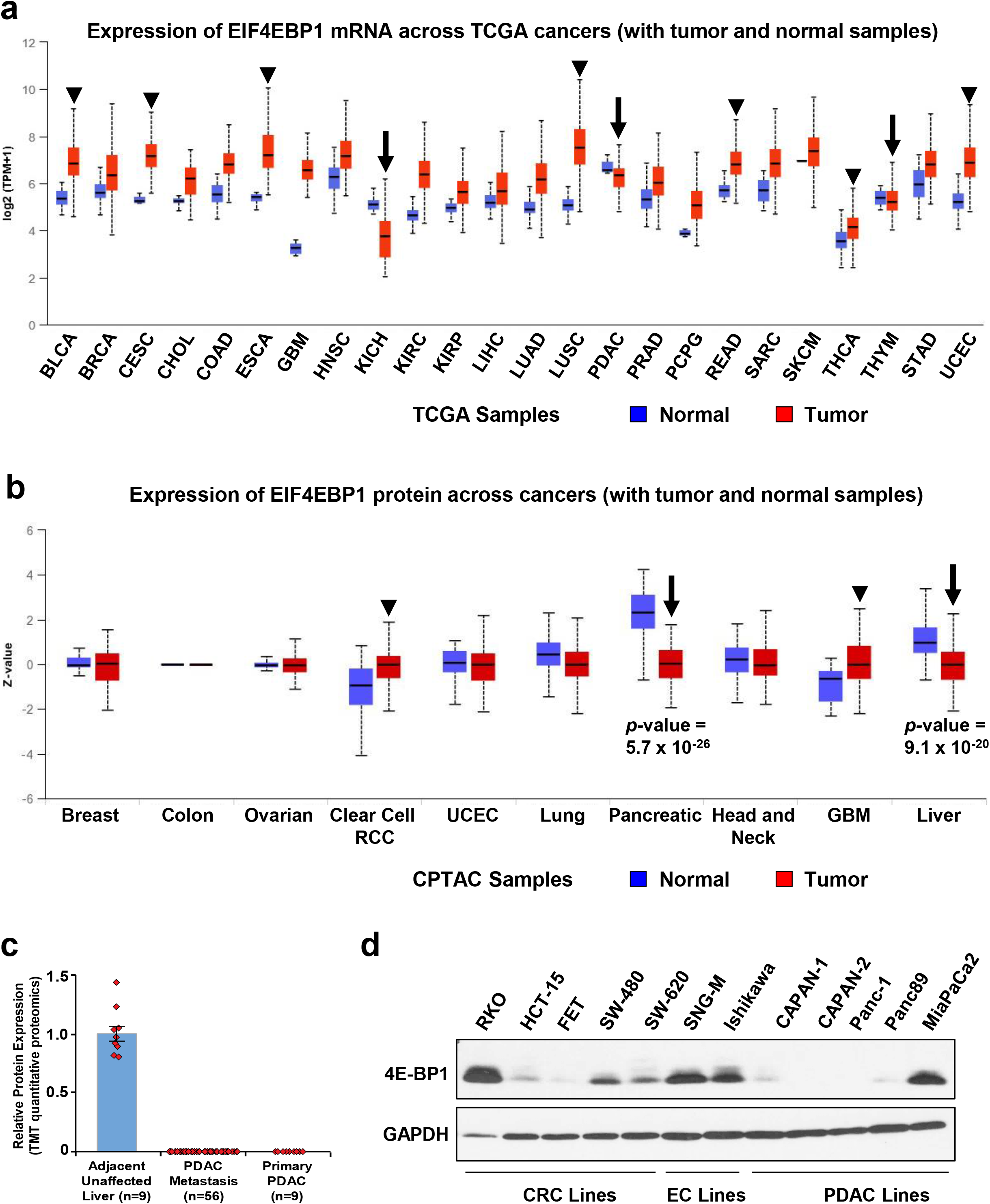
4E-BP1 expression in tumors. ***a***. TCGA RNAseq data on the expression of *EIF4EBP1* (*4E-BP1*) mRNA in the indicated normal and tumor tissues generated using the UALCAN portal. ***b***. As in *A* except that the relative expression of 4E-BP1 protein derived from CPTAC data is shown. The arrows and arrowheads in *A* and *B* indicate tumor types in which 4E-BP1 is downregulated or upregulated relative to normal tissue, respectively. ***c***. LC-MS/MS analysis of the expression of 4E-BP1 protein in primary PDAC, PDAC liver metastasis and adjacent normal liver. ***d***. Western blot analysis of 4E-BP1 and GAPDH expression in the indicated cells lines. CRC: colorectal cancer: EC: endometrial cancer; PDAC: pancreatic adenocarcinoma. Data in *d* are representative of more than 3 independent experiments.

Evidence from our group and others supports a role for PP2A as a 4E-BP1 phosphatase (22–24), although the regulatory B subunits responsible for 4E-BP1 activation have yet to be identified (47). As discussed above, SMAPs target a limited subset of PP2A heterotrimers containing B56α, B56β, or B56ε regulatory subunits. Thus, we used lead SMAP, DT-061, to examine the ability of B56α-, B56β-, and/or B56ε-PP2A to activate 4E-BP1 by dephosphorylation. Initial analysis was performed in IEC-18 rat intestinal crypt-like cells, which were used by our group to demonstrate the ability of PP2A to mediate hypophosphorylation and activation of 4E-BP1 by PKCα signaling (22). As shown in Fig. 2a(i), 4E-BP1 phosphoforms can be readily distinguished by Western blotting in these cells: the slower migrating γ form represents inactive hyperphosphorylated 4E-BP1 and the β/α forms indicate translation repressive hypophosphorylated species (48). The SMAP DT-061 led to rapid accumulation of faster migrating species of 4E-BP1 in these cells (Fig. 2a(ii); *upper panel*), with robust accumulation of the active β form observed by 15 min of treatment. By 15-30 min, the α form was also readily detectable. Western blot analysis further demonstrated that DT-061 promotes loss of phosphorylation at Ser64 and Thr45 (numbering for murine 4E-BP1), as shown using antibodies specific for phosphorylated Ser64 and non-phosphorylated Thr45 residues, respectively (Fig. 2b). Note that only fully phosphorylated 4E-BP1 is incapable of interacting with eIF4E and loss of phosphorylation at Ser64/65 is sufficient for eIF4E binding (8). SMAP-induced accumulation of faster migrating species of 4E-BP1 was also seen in human cancer cells. Optimization of our immunoblotting technique for detection of 4E-BP1 phosphoforms from human cells, which are more difficult to resolve using SDS PAGE, revealed a robust mobility shift in 4E-BP1 in low (Capan-1) and high (Ishikawa) expressors (*cf* Fig. 1d) by 1 h of treatment (Fig. 2c(i)).

**Figure 2.**
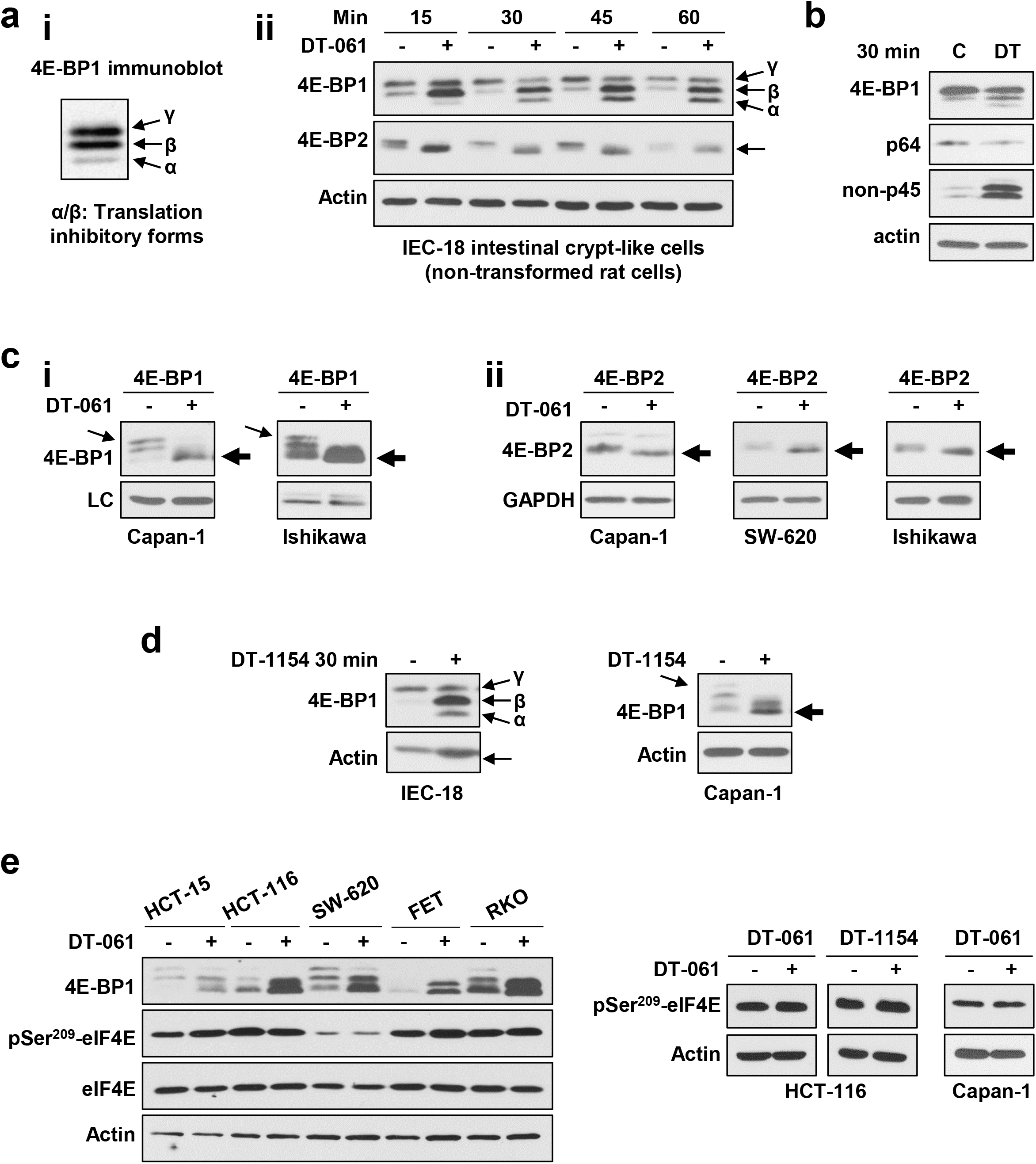
SMAP treatment leads to hypophosphorylation of 4E-BP1 and 4E-BP2. ***a***.***i***. Detail of a Western blot of 4E-BP1 from IEC-18 cells showing the α, β, and γ phosphoforms. ***ii***. IEC-18 cells were treated with 20 µM SMAP DT-061 (+) or vehicle (-) for 15 to 60 min and expression of the indicated proteins was determined by Western blotting. 4E-BP1 phosphoforms and hypophosphorylated 4E-BP2 are indicated with arrows. ***b***. As in *a*, except that cells were treated with DT-061 or vehicle for 30 min and Western blots were probed with antibodies recognizing total protein or specific for 4E-BP1 phosphorylated at Ser64 (p64) or lacking phosphorylation at Thr45 (non-p45). ***c***. Tumor cells were treated with vehicle or 20 µM DT-061 prior to Western blotting for 4E-BP1 (***i***) or 4E-BP2 (***ii***). LC: non-specific band used as a loading control to confirm even loading. Arrows indicate hypophosphorylated 4E-BP1 and 4E-BP2. ***d.*** As in ***a*** and ***c***, except that cells were treated with SMAP DT-1154. ***e.*** Cells were treated with DT-061, DT-1154 or vehicle (-) for 3 h and analyzed for the expression of the indicated proteins by Western blotting. All data are representative of 3 or more independent experiments.

DT-061 also promoted rapid accumulation of faster migrating forms of 4E-BP2 in IEC-18 cells and tumor cells (Fig. 2a(ii),c(ii); arrows), indicating that both 4E-BP1 and 4E-BP2 can be targeted with SMAPs. Importantly, the effects of DT-061 on 4E-BP1 and 4E-BP2 were recapitulated with DT-1154, another well characterized compound in the SMAP series (49) (Fig. 2d and data not shown). In sharp contrast to 4E-BP1 and 4E-BP2, 4E-BP3 is only weakly phosphorylated and appears to be predominantly regulated by transcriptional induction (50). Furthermore, as noted by others (50), 4E-BP3 was not detected in most of the models used in this study (e.g., Fig. S1). Since 4E-BP3 could not be detected either in the absence or presence of SMAP, we were unable to determine if SMAP treatment affects phosphorylation of 4E-BP3. Together, the data point to the ability of the SMAP targets, B56α-, B56β, and/or B56ε-PP2A, to modulate the phosphorylation of 4E-BP1 and 4E-BP2.

PP2A is known to regulate the phosphorylation and activity of other components of the translation machinery, including eIF4E and the eIF4E kinases, Mnk1 and Mnk2 (51). Mnk1/2 phosphorylate eIF4E at Ser^209^ to enhance eIF4E cap binding activity, and PP2A has been shown to negatively regulate eIF4E Ser^209^ phosphorylation through direct dephosphorylation of Mnk1 and eIF4E. However, in contrast to their effects on 4E-BP1 phosphorylation, SMAPs DT-061 and DT-1154 failed to reduce phosphorylation of eIF4E at Ser^209^ in multiple cell lines (Fig. 2e), pointing to an inability of these agents to activate eIF4E- and Mnk-targeting PP2A heterotrimer(s) and highlighting the selectivity of SMAP compounds.

Treatment of cells with SMAP DT-061 for longer times (e.g., 3 h) revealed an unexpected ability of SMAPs to markedly upregulate 4E-BP1 protein (Fig. 3a,b). The effect, which was concentration-dependent and sustained for at least 72 h (Fig. 3a,b), was observed in every PDAC, EC, and CRC cell line tested (Fig. 3c-e). Notably, SMAPs restored the expression of 4E-BP1 in cells with profound repression of the protein (e.g., many PDAC cell lines, Fig. 3c) and increased the expression of 4E-BP1 in cells expressing readily detectable basal levels of the protein (e.g., MiaPaCa-2, S2-013, SNG-M, Fig. 3c,d). Upregulation of 4E-BP1 was also seen with SMAP DT-1154 (Fig. 3f) and was noted in SMAP-treated patient-derived colorectal and pancreatic cancer organoids (Fig. 3g), models that closely recapitulate the tumors from which they were derived (52). The effect appears to be specific for 4E-BP1 since levels of 4E-BP2 (Fig. 3h, *upper panel*) and 4E-BP3 (Fig. 3i) were not affected by prolonged SMAP treatment. In the case of 4E-BP3, Rapamycin (Rap) was used as a positive control for induction of the protein in MiaPaCa cells (as previously described (50)). Since many cancers overwhelm the inhibitory capacity of cellular 4E-BP1 by upregulating eIF4E to decrease the 4E-BP1:eIF4E ratio (5,16,17), the finding that SMAPs can increase basal levels of 4E-BP1 protein points to the promise of these agents for restoring translational control in a broad spectrum of tumors.

**Figure 3.**
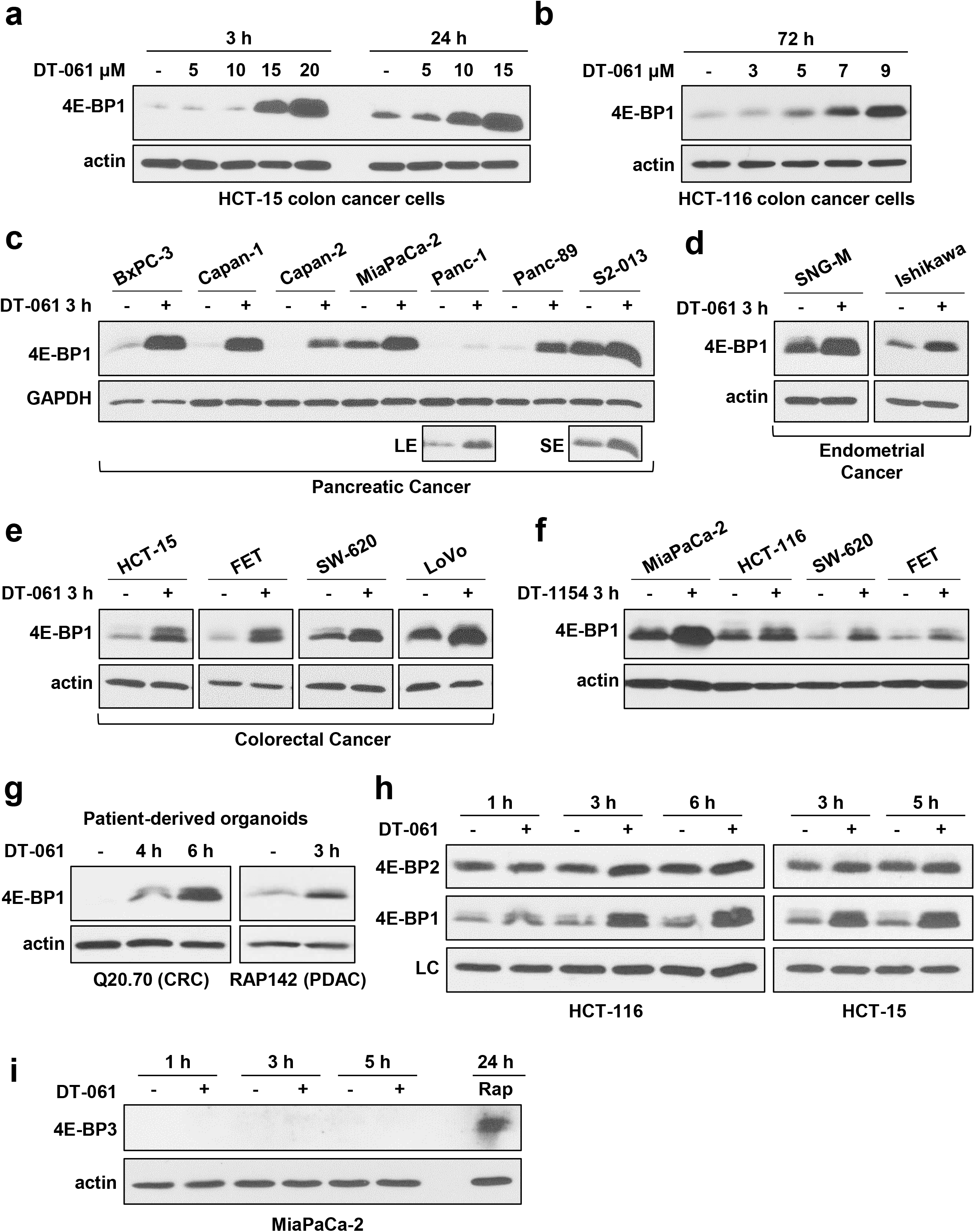
SMAP treatment upregulates 4E-BP1 but not 4E-BP2 or 4E-BP3. ***a-e***. CRC, PDAC, or EC cell lines were treated with vehicle or 20 µM DT-061 for 3 h, 24 h, or 72 h before analysis by Western blotting for the indicated proteins. LE: long exposure: SE: short exposure. ***f.*** The indicated cell lines were treated with 20 µM SMAP DT-1154 for 3 h and subjected to anti-4E-BP1 or -actin immunoblotting. ***g.*** Patient-derived CRC or PDAC organoids were treated with DT-061 for the indicated times and analyzed as in *f*. ***h.*** Cells were treated with DT-061 for the indicated times and expression of 4E-BP2 and 4E-BP1 was assessed by Western blotting. LC: non-specific band used as a loading control to confirm even loading. ***i.*** Cells were treated with DT-061 or Rapamycin (Rap), a known inducer of 4E-BP3 expression, for the indicated times and analyzed for 4E-BP3 expression by Western blotting. All data are representative of 3 or more independent experiments.

Having discovered that SMAPs can upregulate 4E-BP1 in tumor cells, we sought to determine if the accumulated 4E-BP1 protein is also hypophosphorylated with SMAP treatment. As shown in Fig. 4a, immunoblot analysis revealed that 4E-BP1 is exclusively in the hypophosphorylated α/β forms at 3 h of SMAP treatment (arrows). Loss of phosphorylation was confirmed in multiple cell types using phospho-Ser65-specific and non-phospho-Thr46-specific antibodies (numbering for human 4E-BP1) (Fig. 4b). Analysis of phosphoproteomic data in DT-061-treated H358 lung cancer cells revealed that, in addition to targeting Ser65 and Thr46, SMAPs also promote loss of phosphorylation at Thr70 and Thr36/37 (Fig. 4c). Together, the data demonstrate that SMAPs promote upregulation of 4E-BP1 protein by 3 h of treatment as well as rapid loss of phosphorylation at all major phosphosites, including Thr37, Thr46, Thr70 and Ser65 (Fig. 4d).

**Figure 4.**
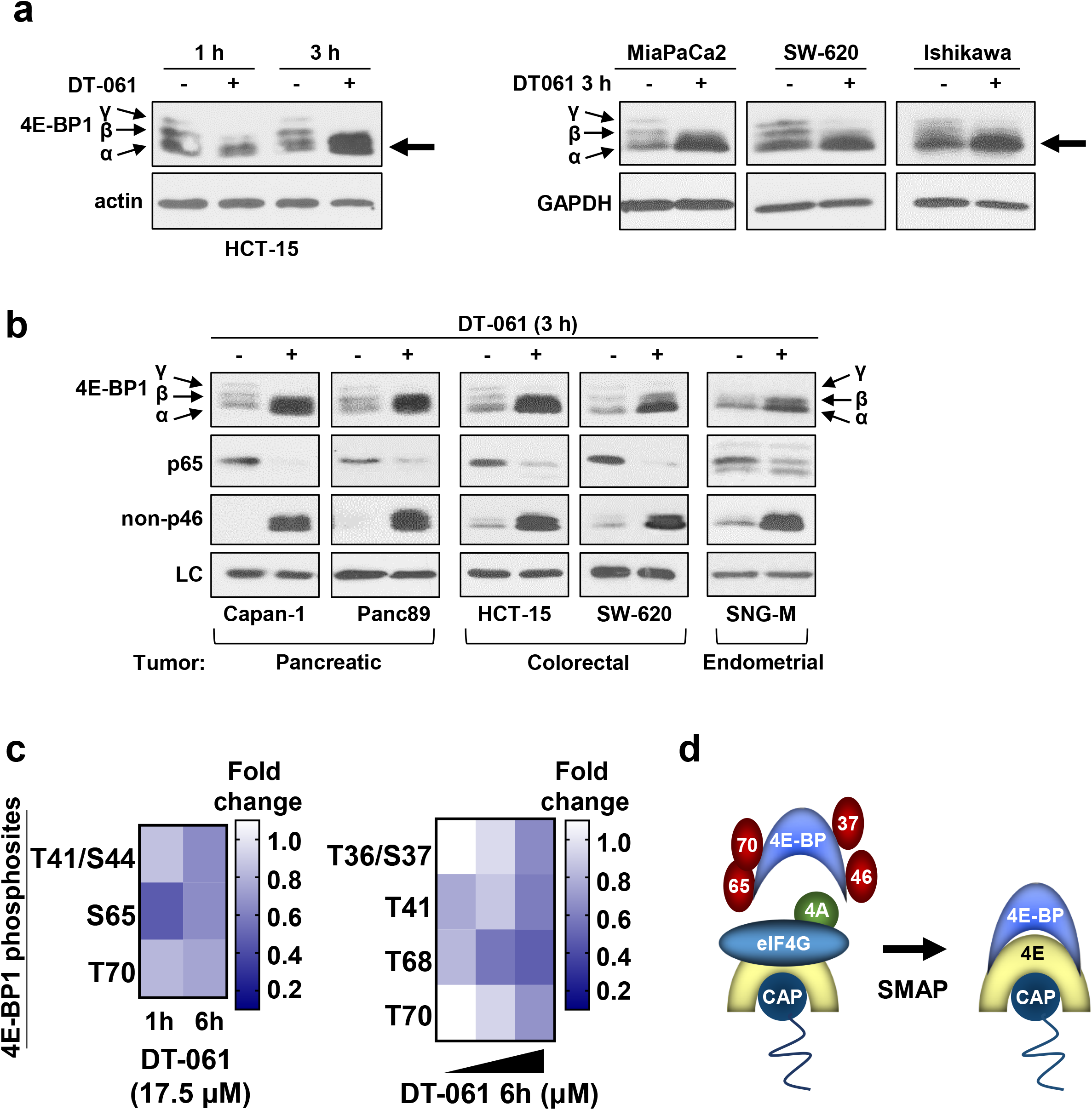
SMAP treatment leads to the accumulation of hypophosphorylated 4E-BP1 in tumor cells. ***a,b***. The indicated cell lines were treated with vehicle or 20 µM DT-061 for 1 h or 3 h before expression and electrophoretic mobility of the indicated proteins was determined by Western blotting. LC: actin or GAPDH used as a loading control. Data are representative of at least three independent experiments. ***c***. Phosphoproteomic analysis reveals that 4E-BP1 is hypophosphorylated upon DT-061 treatment in H358 lung cancer cells. LC-MS/MS analysis of 4E-BP1 phosphorylation in cells treated with 17.5 µM DT-061 for 1 h or 6 h (*left panel*) or with increasing doses (17.5, 22.5, and 30 µM DT-061) for 6 h (*right panel*). Heatmaps show the average of 3 biological replicates. ***d***. Model showing effects of SMAP treatment on 4E-BP phosphorylation and eIF4E (4E) binding, with displacement of eIF4G and eIF4A (4A).

### SMAP-induced 4E-BP1 upregulation and dephosphorylation are B56-PP2A-dependent

The PP2A dependence of SMAP-induced effects on 4E-BP1 was supported by the ability of two compounds from the SMAP series, DT-061 and DT-1154, to modulate the expression and phosphorylation of the protein (Figs. 1-3). Several additional approaches were used to conclusively establish the PP2A dependence of SMAP effects. First, we used DT-1310 and DT-766 (38,53), compounds that are structurally similar to DT-061 but unable to stabilize/activate PP2A (for structures, see Supplementary Data, Fig. S2). While DT-1310 does not bind to PP2A, DT-766 binds to the A/C dimer but does not stabilize its association with B subunits. DT-766 thus acts as a competitive inhibitor of the PP2A-dependent effects of DT-061. Neither DT-1310 nor DT-766 induced upregulation or hypophosphorylation of 4E-BP1 (Fig. 5a,b), and a 4-fold excess of DT-766 blocked the effects of DT-061 (Fig. 5b). c-myc, a known target of SMAPs, was used as a positive control for the effects of SMAPs and DT-766 (Fig. 5b). PP2A dependence was also confirmed by the ability of pharmacological inhibitors of PP2A, okadaic acid (OA) and calyculin A (Cal-A), to block the effects of SMAP treatment in a panel of PDAC and CRC cell lines (Fig. 5c).

**Figure 5.**
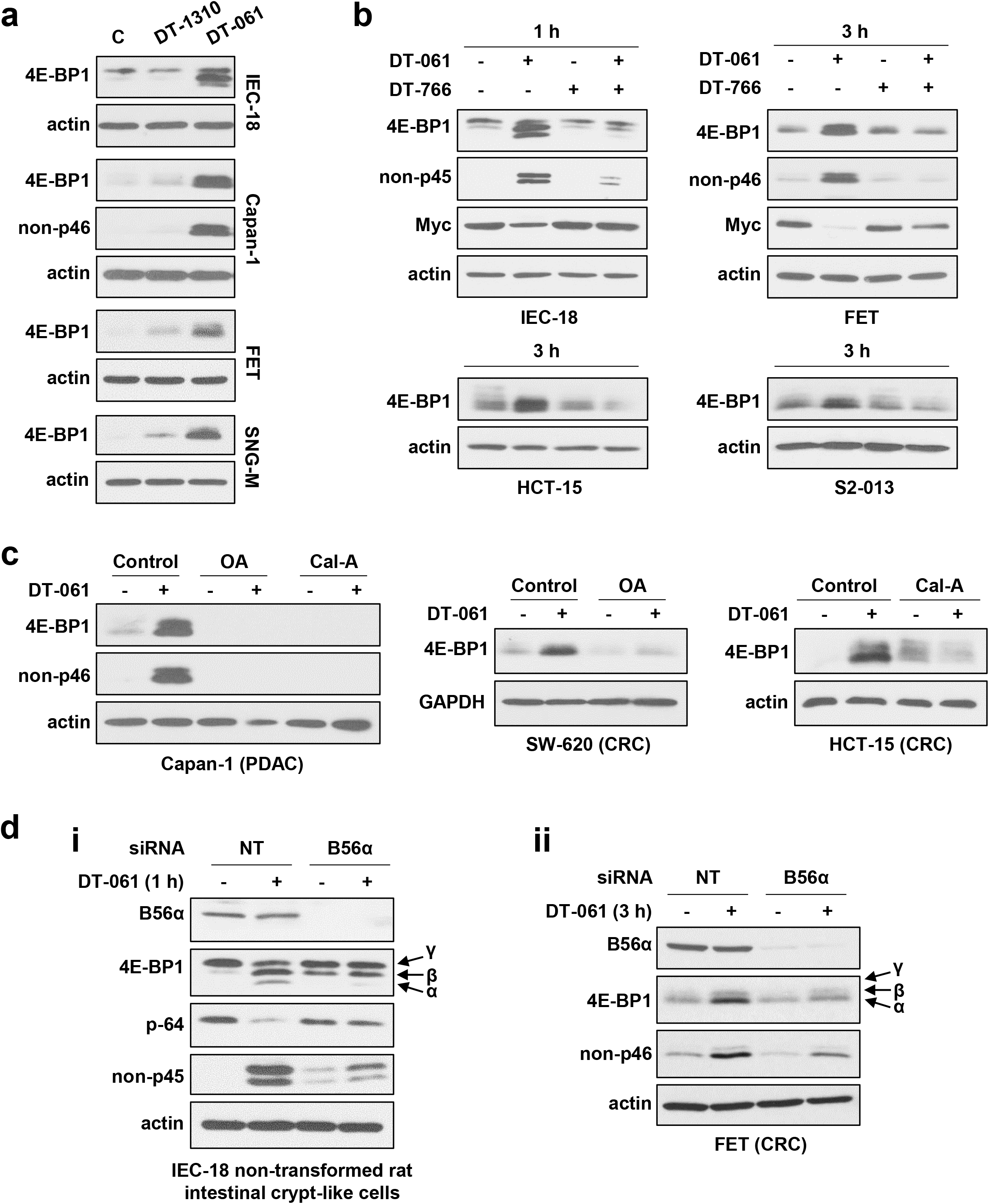
Effects of SMAPs on 4E-BP1 are mediated by PP2A. ***a***. The indicated cell lines were treated with vehicle, 20 µM DT-1310 or 20 µM DT-061 and 4E-BP1 and actin levels were detected by Western blotting. Changes in 4E-BP1 phosphorylation were detected using an antibody that detects 4E-BP1 only when lacking phosphorylation at Thr45/46 (non-p45/p46). IEC-18 cells and human tumor cells (Capan-1, FET, SNG-M) were treated for 1 h and 3 h, respectively. ***b***. As in *a* except that cells were treated with 20 µM DT-061 and 80 µM DT-766 alone or in combination. ***c***. The indicated cell lines were pretreated with 120 nM okadaic acid (OA) or 6 nM calyculin A (Cal-A) prior to the addition of 20 µM DT-061 (SMAP) for 3 h. Expression of the indicated proteins was analyzed by Western blotting as in *a*. ***d***. Cells were transfected with non-targeting (NT) or ONTARGETplus SMARTpool siRNA against B56α (PPP2R5A) for 48 h (IEC-18) or 48 h + 72 h (FET), respectively, before treatment with vehicle (C) or 20 µM DT-061 (DT) for 1 or 3 h as indicated. Expression, migration and phosphorylation of the indicated proteins was then determined by Western blotting. p-64: phospho-Ser64 of 4E-BP1; non-p45/non-p46: 4E-BP1 lacking phosphorylation at Thr45/46. All data are representative of 3 or more independent experiments.

While the SMAP-binding pocket accommodates B56α, B56β and B56ε, our previous analysis indicates that B56α may be the major regulatory subunit targeted by SMAPs (37, and unpublished data); therefore, the role of this subunit in the effects of SMAPs was tested using siRNA. As shown in Fig. 5d(i,ii), B56α knockdown inhibited DT-061-induced 4E-BP1 hypophosphorylation in IEC-18 cells and FET CRC cells, as confirmed using phospho-specific antibodies. B56α knockdown also inhibited SMAP-induced increased expression of 4E-BP1 (Fig. 5d**ii)**. These findings establish a role of B56α-PP2A in regulation of 4E-BP1 expression and phosphorylation. However, B56α knockdown did not completely block the effects of SMAP, indicating that additional SMAP targets, B56β-PP2A and/or B56ε-PP2A, may also function to upregulate and dephosphorylate 4E-BP1.

### SMAP-induced hypophosphorylation of 4E-BP1 is not the result of inhibition of upstream regulators

The ability of SMAP/B56-PP2A to negatively regulate phosphorylation at all major 4E-BP1 phosphosites could reflect inhibition of upstream kinase(s) or direct dephosphorylation. A major upstream regulator of 4E-BP1 is mTOR, which can directly phosphorylate Thr37, Thr46, Thr70 and Ser65 to inactivate the protein (Fig. 6a). The activity of mTOR, in turn, is regulated by upstream AKT and ERK signaling and ERK has been implicated in direct phosphorylation of Ser65 on 4E-BP1 (8,14,15) (Fig. 6a). Analysis of activating phosphorylation of AKT at Ser473 and Thr308 excluded effects of SMAP treatment on upstream AKT activation, consistent with evidence that B55α-PP2A, but not B56-PP2A, dephosphorylates AKT (54) (Fig. 6b). Although DT-061 reduced ERK phosphorylation/activity in some cell lines (e.g., Panc-1, Panc-89), ERK phosphorylation was unaffected in a majority of cell lines tested (Fig. 6c), indicating that SMAP effects on 4E-BP1 are unlikely to be mediated by alterations in ERK activity. To test the possibility that SMAPs hypophosphorylate 4E-BP1 by inhibiting mTOR, we examined phosphorylation of the mTOR substrates S6 kinase and ULK1. As shown in Fig. 6d(i,ii), DT-061 did not induce changes in phosphorylation of S6 kinase or ULK1 in multiple cell types, indicating that the effects of SMAPs on 4E-BP phosphorylation are not mediated by inhibition of mTOR. In contrast, the mTOR inhibitor, INK128, readily inhibited the phosphorylation of mTOR substrates such as ULK1 (Fig. 6dii, *left panel*). The finding that DT-061 hypophosphorylates 4E-BP1 in cells with functional AKT, ERK, and mTOR supports the conclusion that SMAP/B56-PP2A effects are not mediated by inhibition of upstream kinase(s). Thus, changes in 4E-BP1 phosphorylation likely reflect direct dephosphorylation of 4E-BP1 protein by B56α-PP2A, and possibly B56β-PP2A and/or B56ε-PP2A.

**Figure 6.**
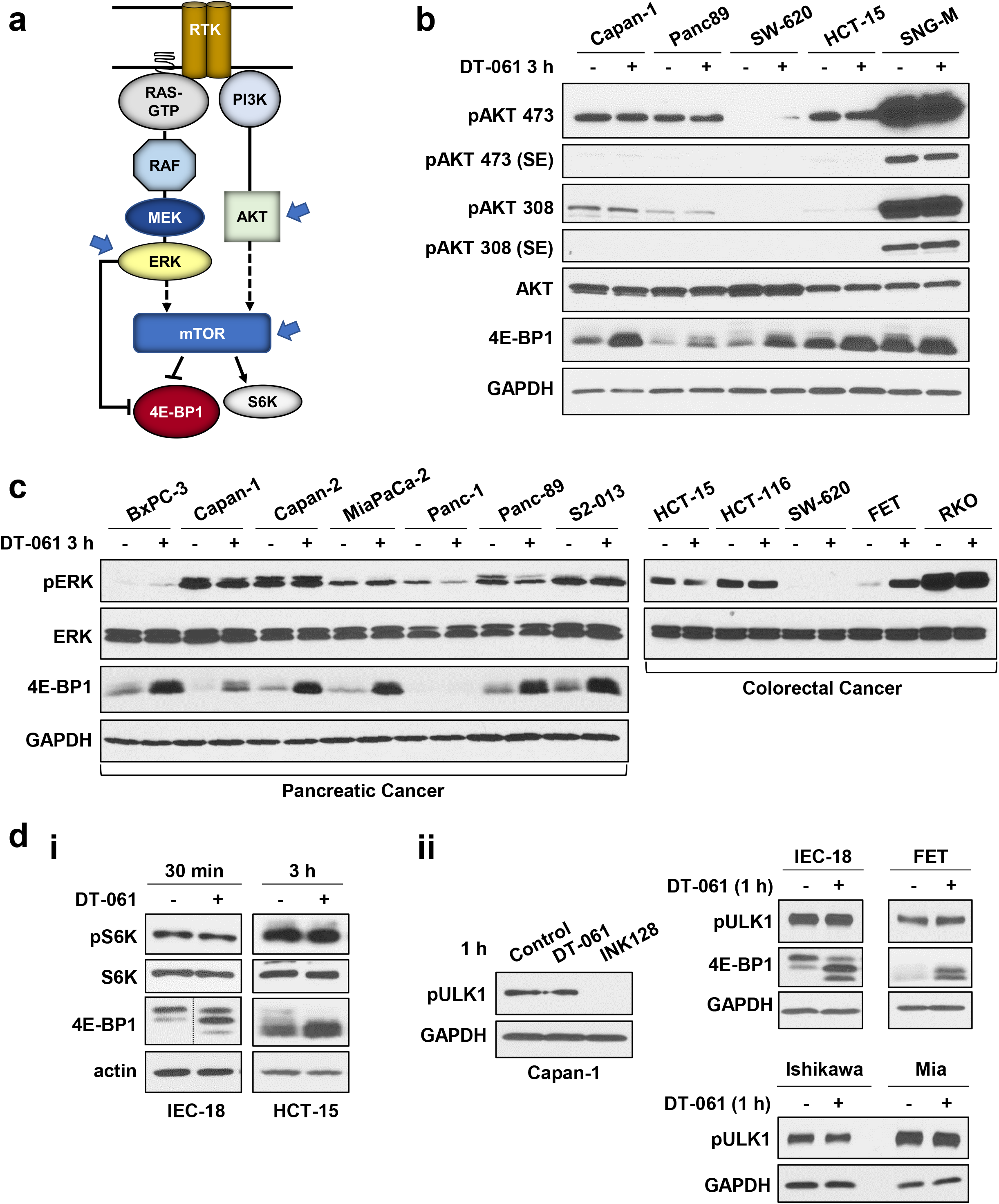
Inhibition of upstream kinases does not account for effects of SMAP/B56-PP2A on 4E-BP1. ***a***. Model showing upstream kinases that regulate phosphorylation of 4E-BP1. mTOR is a major 4E-BP1 kinase and ERK has also been implicated in phosphorylation of specific sites on the protein. AKT and ERK are also upstream regulators of mTOR activity. ***b-d***. The indicated cell lines were treated with vehicle or 20 µM DT-061 for 30 min, 1 h or 3 h before analysis of expression of total AKT, ERK1/2 (ERK), S6K, 4E-BP1 and GAPDH as well as phosphorylation of AKT (pAKT) on Ser473 and Thr308, ERK1/2 on Thr202/Thr204 (pERK), S6 kinase on Thr389 (pS6K), and ULK1 on Ser757 (pULK1) by Western blotting. INK128: mTOR inhibitor used to demonstrate that mTOR inhibition abrogates phosphorylation of ULK1 on Ser757. Note that the CRC samples analyzed in *c* are the same as in *2e*; see Fig. *2e* for confirmation of effects on 4E-BP1. All data are representative of 3 or more biological replicates.

### SMAP/B56-PP2A activates the translation repressive functions of 4E-BP1 in multiple cell types

Three approaches were used to evaluate the functional consequences of SMAP/B56-PP2A-induced upregulation and dephosphorylation of 4E-BP1: cap affinity assays, eIF4E immunoprecipitation (eIF4E co-IP) assays, and bicistronic reporter cap-dependent translation activity assays (55,56). While cap affinity assays use m^7^GTP cap analog-conjugated agarose beads to mimic the 5’-mRNA cap and capture eIF4E and its binding partners, eIF4E co-IP assays use anti-eIF4E antibodies to isolate eIF4E for analysis (Fig. 7a). Thus, these assays reveal changes in the interaction of 4E-BP1 with eIF4E. Immunoblot analysis of cap analog-bound proteins revealed a striking increase in levels of cap-bound 4E-BP1 in DT-061-treated cells, with concomitant displacement of eIF4G, in multiple tumor cell types (Fig. 7b and data not shown). A similar increase was seen in cap-bound 4E-BP2 (Fig. 7b, *left panel*). eIF4E co-IP assays confirmed these findings, with DT-061 inducing a marked increase in 4E-BP1-eIF4E and 4E-BP2-eIF4E interaction, accompanied by a decrease in eIF4E-bound eIF4G in CRC, EC, and PDAC cell lines (Fig. 7c and data not shown). Together, these findings confirm that B56-PP2A activation by SMAPs inhibits the formation of the eIF4F translation initiation complex in multiple cancer cell types.

**Figure 7.**
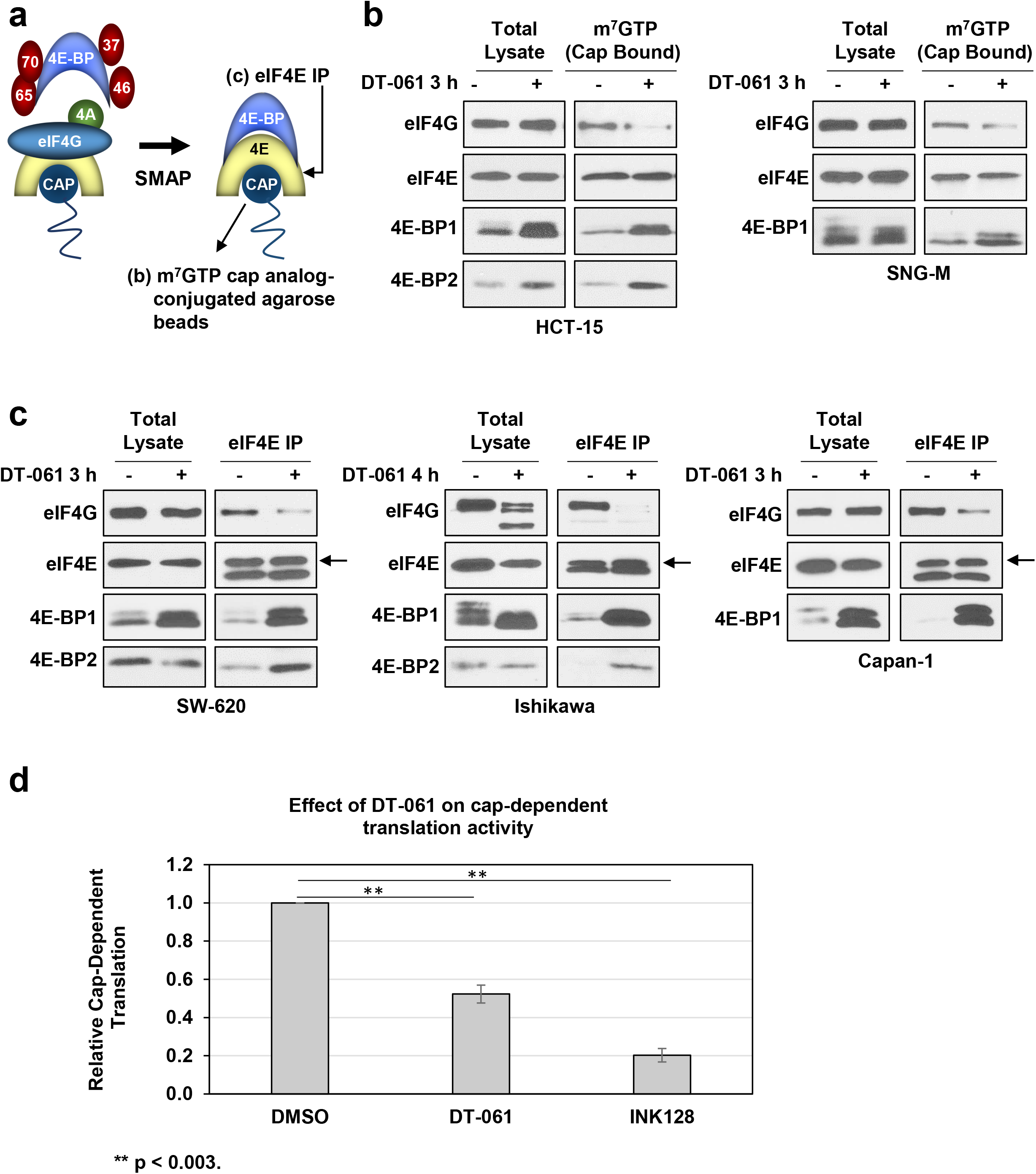
SMAP/B56-PP2A functionally activates 4E-BPs. ***a***. Effects of SMAP treatment on the interaction between eIF4E and 4E-BP1 were assessed by m^7^GTP cap pulldown assays (see *b*) and eIF4E IP assays (see *c). **b.*** HCT-15 and SNG-M cells were treated with vehicle or 20 µM DT-061 for the indicated times. Cells were then lysed and cap-binding proteins were pulled down using m^7^GTP-Agarose. Levels of the indicated proteins in m^7^GTP cap pulldowns (Cap-Bound) and the original lysates (Total Lysate) were determined by Western blotting. ***c***. As in *b* except that eIF4E was immunoprecipitated from lysates (eIF4E IP) before analysis of the original lysates and co-immunoprecipitated proteins by Western blotting. Arrows indicate eIF4E above immunoglobulin light chain. ***d***. Capan-1 cells were transfected with pcDNA3-RLUC-POLIRES-FLUC dual luciferase reporter plasmid before treatment with vehicle, 20 µM DT-061 (DT) or 10 nM INK128. After 6 h, firefly and Renilla luciferase activity was determined, and the ratio of cap-dependent Renilla luciferase expression and IRES-driven firefly luciferase expression was calculated and expressed as relative to the ratio in vehicle control cells. Data for each cell line in *b* and *c* are representative of at least 2 independent experiments. Data in *d* are the average of 4 independent experiments. ** : p < 0.003.

Follow-up experiments assessed the ability of DT-061 to suppress cap-dependent translation using a bicistronic dual luciferase reporter system that directly measures cap-dependent translation activity by monitoring the ratio between cap-dependent and cap-independent translation (55,56). As shown in Fig. 7d, this analysis confirmed that SMAP suppresses cap-dependent translation (Fig. 7d). Together, the data from cap affinity assays, eIF4E co-IP assays and cap-dependent translation activity assays show that B56-PP2A activation is able to counter all mechanisms used by tumor cells to disable 4E-BP1 (i.e., hyperphosphorylation of 4E-BP1, downregulation of 4E-BP1, and increased eIF4E:4E-BP1 ratio) and thereby promote 4E-BP1-mediated inhibition of cap-dependent translation in a variety of cancer cell types.

### SMAP/B56-PP2A transcriptionally activates the 4E-BP1 gene (EIFEBP1)

Next, we explored the mechanisms downstream of SMAP-induced B56-PP2A activation that mediate upregulation of 4E-BP1. Time course analysis revealed that SMAP/PP2A-induced upregulation of the protein is first detected by 2 to 3 h of treatment dependent on the cell type (Fig. 8a). RT-qPCR analysis further showed that SMAP treatment induces 4E-BP1 mRNA with a similar time course (Fig. 8b). Upregulation of 4E-BP1 protein was blocked by the transcription inhibitor actinomycin D (ActD), pointing to a transcriptional mechanism (Fig. 8c). The ability of DT-061 to enhance 4E-BP1 mRNA synthesis rates was further confirmed using the Invitrogen ethylene uridine (EU)-based Click-IT Nascent RNA Capture Kit, which determined that B56-PP2A activation promotes an ∼4-fold increase in 4E-BP1 mRNA synthesis by 6 h in Capan-1 cells (Fig. 8c). Thus, B56-PP2A activation transcriptionally activates the *EIF4EBP1* gene to promote the accumulation of the translational repressor in multiple cancer types.

**Figure 8.**
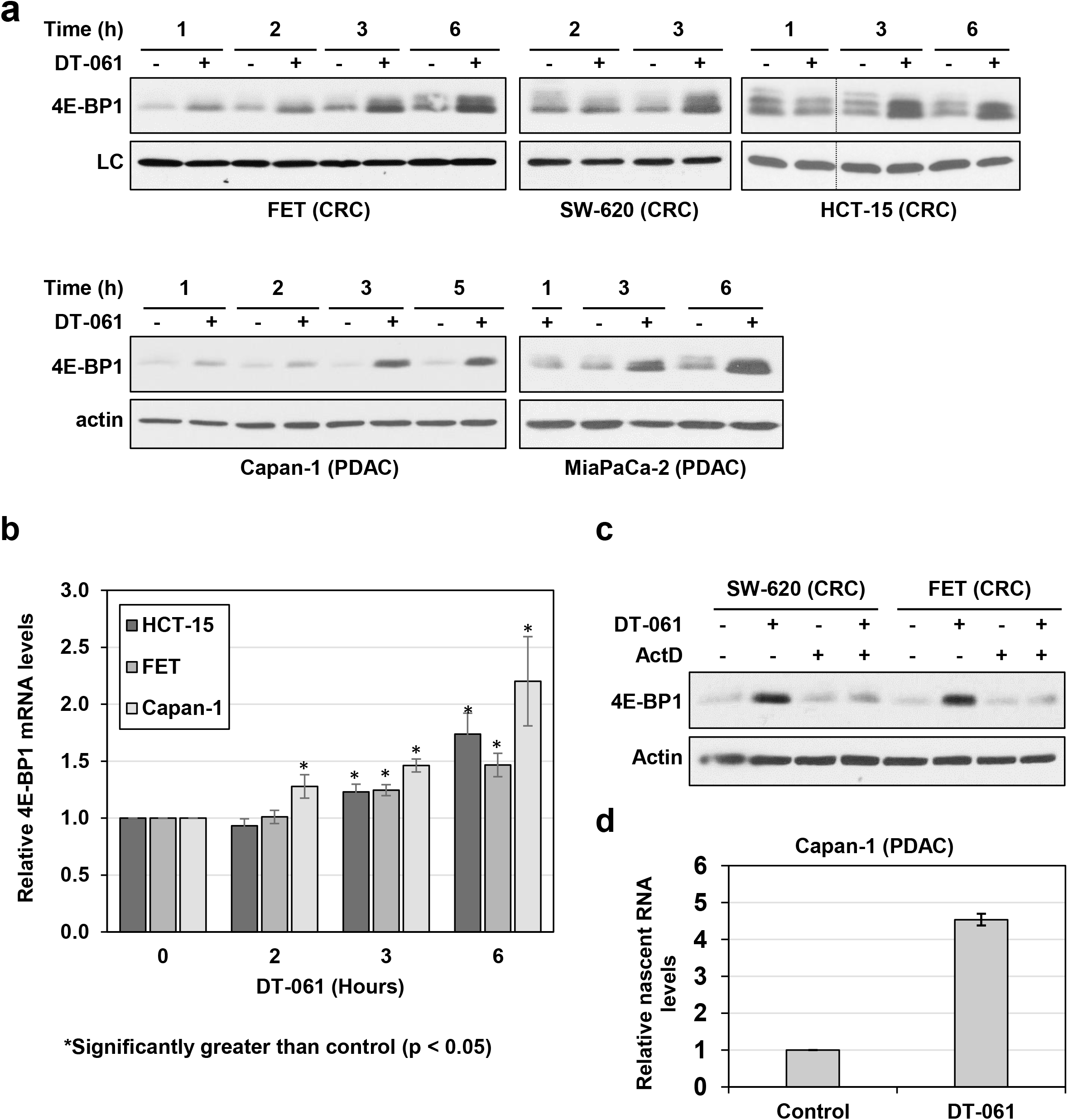
SMAP/B56-PP2A induces upregulation of 4E-BP1 at the level of transcription. ***a***. Cells were treated with 20 µM DT-061 as indicated and expression of 4E-BP1 and actin/GAPDH loading control (LC) was determined by Western blotting. Dashed lines indicate rearrangement of lanes from a single blot for clarity. ***b***. As in *a* except that 4E-BP1 mRNA expression was determined by RT-qPCR, normalized to 18 S rRNA levels and expressed as relative to vehicle control. ***c.*** Cells were pretreated with 1 μg/ml actinomycin D (ActD) for 1 h as indicated prior to addition of vehicle or DT-061 for 4 h. ***d***. Capan-1 cells were treated with vehicle or 20 µM DT-061 for 6 h and EU was added during the last hour. Total RNA and EU-labeled RNA (newly synthesized RNA) were then isolated, and 4E-BP1 mRNA was quantified by RT-qPCR, normalized to 18 S rRNA and expressed relative to levels in vehicle-treated cells. Data in *a* and *c* are representative of at least 3 biological replicates for each cell line and data in *b* are the average of at least 3 independent experiments. Data in *d* are the average of 2 independent experiments. * : significantly greater than control (*, p < 0.05).

### Induction of 4E-BP1 gene expression requires ATF4

Mechanistic understanding of the transcriptional regulation of 4E-BP1 is limited (56–59). Previous studies support a role for the transcription factors c-Myc (57), ATF4 (58), and HIF1α (59) as activators 4E-BP1 transcription. We excluded c-Myc because it is downregulated by SMAPs in tumor cells (37) and focused our initial analysis on ATF4 based on evidence that ATF4 directly activates 4E-BP1 transcription during *Drosophila* development and aging (60) and in pancreatic β cells during ER stress (58). Western blot analysis revealed that SMAP treatment increases ATF4 protein levels by 1 h (Fig. 9a), thus preceding the induction of 4E-BP1 mRNA and protein that generally occurs after 2 h (Fig. 9a, *cf* Fig. 8a,b). ATF4 protein levels continued to increase over a 6 h SMAP incubation period (Fig. 9a). SMAP-induced upregulation of ATF4 was observed in all cancer cell lines tested and was also detected in patient-derived tumor organoids (Fig. 9b and data not shown). Consistent with SMAP effects on 4E-BP1 protein, DT-061 upregulation of ATF4 was PP2A-dependent, as confirmed in competition experiments with a 4-fold excess of the SMAP competitive inhibitor DT-766 (Fig. 9c) and using the pharmacological inhibitors of PP2A, OA and Cal-A (Fig. 9d).

**Figure 9.**
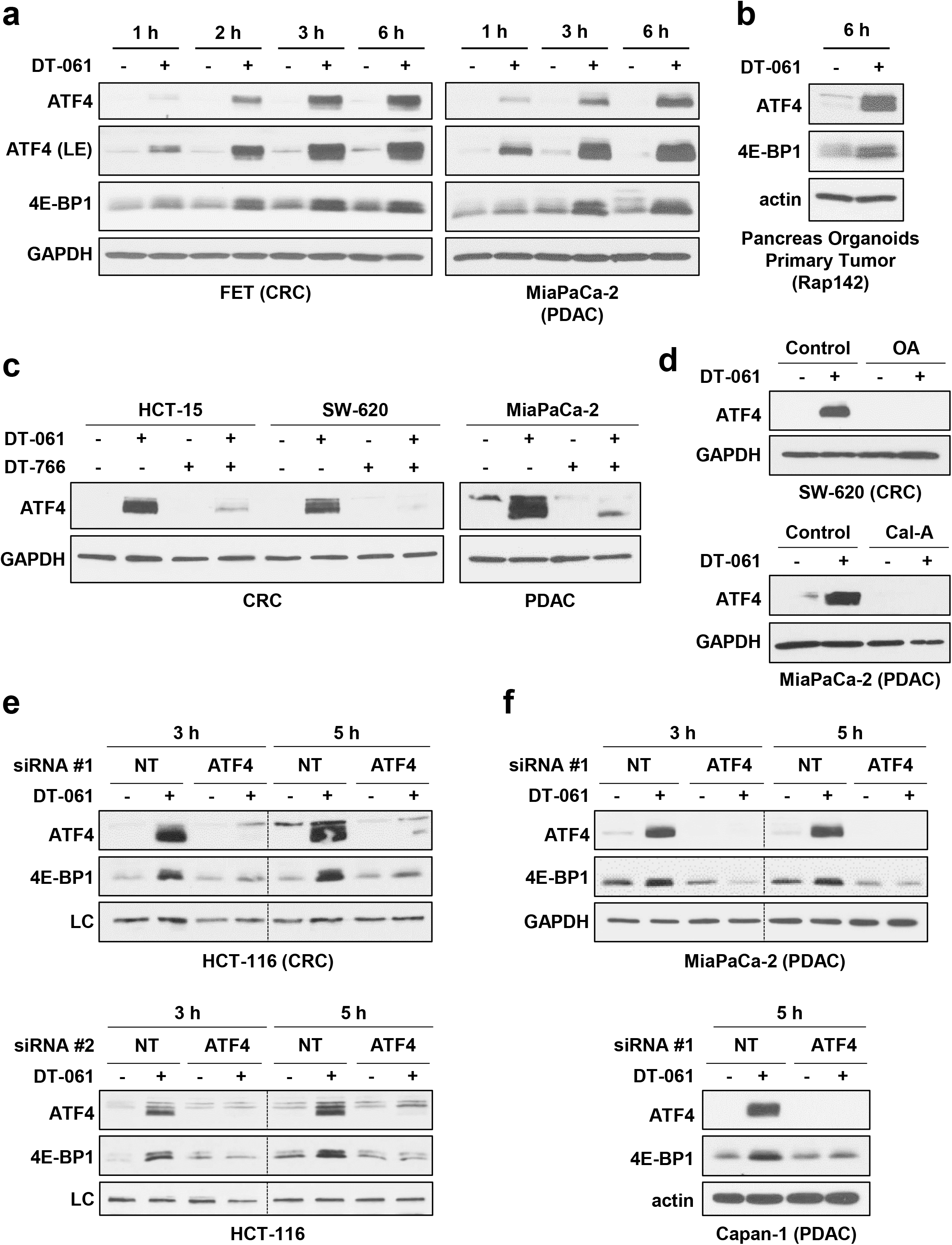
SMAP/B56-PP2A upregulation of 4E-BP1 is dependent on ATF4. ***a,b***. FET cells, MiaPaCa-2 cells or patient-derived pancreatic cancer organoids were treated with vehicle or 20 µM DT-061 for 1 to 6 h and expression of the indicated proteins was analyzed by Western blotting. LE: long exposure. ***c***. As in *a* except that cells were treated with vehicle, 20 µM DT-061 and 80 µM DT-766 alone or in combination. ***d***. The indicated cell lines were pretreated with 120 nM okadaic acid (OA) or 6 nM calyculin A (Cal-A) prior to the addition of 20 µM DT-061 for 3 h. Expression of the indicated proteins was then analyzed by Western blotting. ***e,f.*** HCT-116, MiaPaCa-2, and Capan1 cells were transfected with non-targeting (NT) or ATF4 targeting siRNA (#1 or #2) for 24 h prior to treatment with vehicle or 20 µM DT-061 for the indicated times. Expression of the indicated proteins was then analyzed by Western blotting. LC: non-specific band used as a loading control. Data in *a-d* are representative of 3 or more independent experiments. Data from each cell line in *e* and *f* are representative of at least 2 independent experiments

Next, we tested the requirement for ATF4 in SMAP/B56-PP2A upregulation of 4E-BP1 using siRNA technology. As shown in Fig. 9e, knockdown of ATF4 with two different siRNAs (*Top panel:* siRNA #1; *Bottom panel:* siRNA #2) abrogated SMAP-induced upregulation of 4E-BP1 in HCT-116 CRC cells. This finding was recapitulated in multiple cell lines including MiaPaCa-2 and Capan-1 PDAC cells (Fig. 9f), confirming that ATF4 plays a requisite role in the effect. Together, the data support a model in which activation of B56-PP2A induces the expression of ATF4, and ATF4 in turn activates the 4E-BP1 promoter to increase 4E-BP1 gene expression. This model is consistent with ATF4 Chip-seq data from the ENCODE database showing strong association of ATF4 with a site in intron 1 of the *EIF4EBP1* gene (Fig. S2) and with previous studies in *Drosophila* (60) and pancreatic β cells (58).

### SMAP/B56-PP2A activates the transcription factors, TFE3/TFEB, by dephosphorylation to promote ATF4 and 4E-BP1 gene expression

Follow-up experiments explored the mechanisms underlying SMAP/B56-PP2A induction of ATF4. RT-qPCR analysis showed that SMAP treatment upregulates ATF4 mRNA by 1-2 h (Fig. 10a), and the ability of the transcription inhibitor Act D to block SMAP-induced upregulation of ATF4 protein pointed to a transcriptional mechanism (Fig. 10b). Together, the data indicated that B56-PP2A activation promotes the transcription of ATF4 in a broad range of cell types.

**Figure 10.**
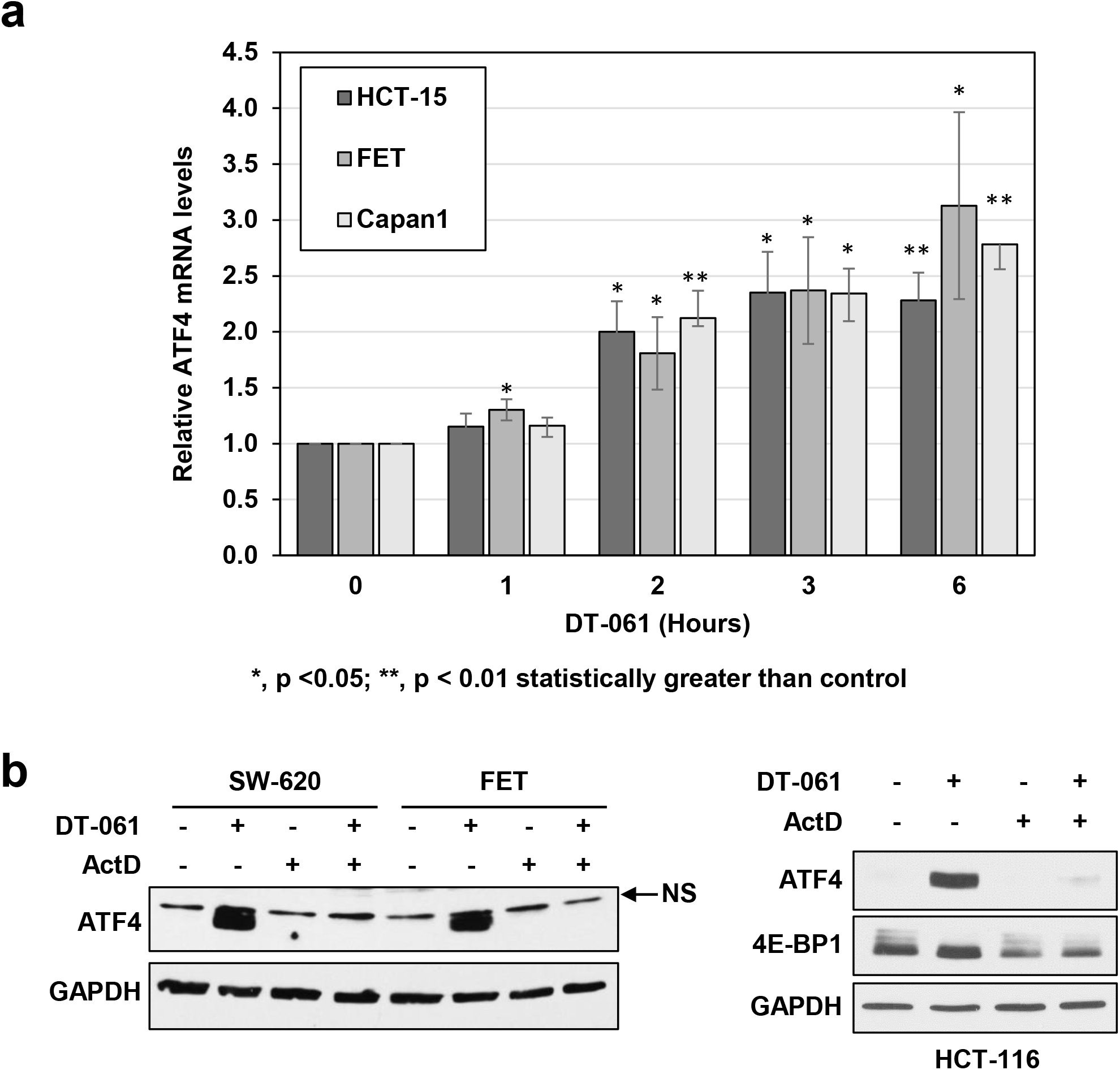
SMAP/B56-PP2A promotes transcriptional upregulation of ATF4. ***a.*** Cells were treated with vehicle or 20 µM DT-061 for the indicated times and ATF4 mRNA was quantified by RT-qPCR, normalized to 18 S rRNA and expressed relative to vehicle-treated controls for each timepoint. ***b***. As in *a* except that cells were pretreated with 1 μg/ml actinomycin D (ActD) as indicated prior to addition of vehicle or DT-061. Data in *a* are the average of at least 3 independent experiments and data in *b* are representative of 3 or more independent experiments. * : p < 0.05, ** p < 0.01 significantly greater than control.

TFE3 and TFEB are two transcription factors that have recently been implicated in regulation of ATF4 gene expression (61). Nuclear accumulation and transcriptional activity of TFE3 and TFEB require dephosphorylation (62,63), with calcineurin (62) and B55α-PP2A (63) emerging as candidate TFE3/TFEB phosphatases. Based on these findings, we tested the ability of SMAPs to regulate the phosphorylation and subcellular localization of these factors. Remarkably, SMAP treatment induced a marked mobility shift of TFE3 (Fig. 11a(i), arrow) and TFEB (Fig. 11a(ii)), characteristic of dephosphorylation and activation of these factors (64), in multiple cell types. The PP2A dependence of SMAP effects on TFE3/TFEB was confirmed using the SMAP competitive inhibitor DT-766 and the PP2A inhibitors, OA and Cal-A (Fig. 11b). Immunofluorescence analysis further confirmed that SMAPs promote rapid nuclear translocation of TFE3/TFEB (Fig. 11c). B55α knockdown did not prevent SMAP-induced dephosphorylation of these factors (Fig. S4), excluding an indirect role of this regulatory PP2A subunit. Thus, the data provide the first indication that B56-PP2A(s) may also be able to dephosphorylate and activate TFE3/TFEB to enhance ATF4 transcription and promote ATF4-dependent transcriptional activation of the *EIF4EBP1* gene.

**Figure 11.**
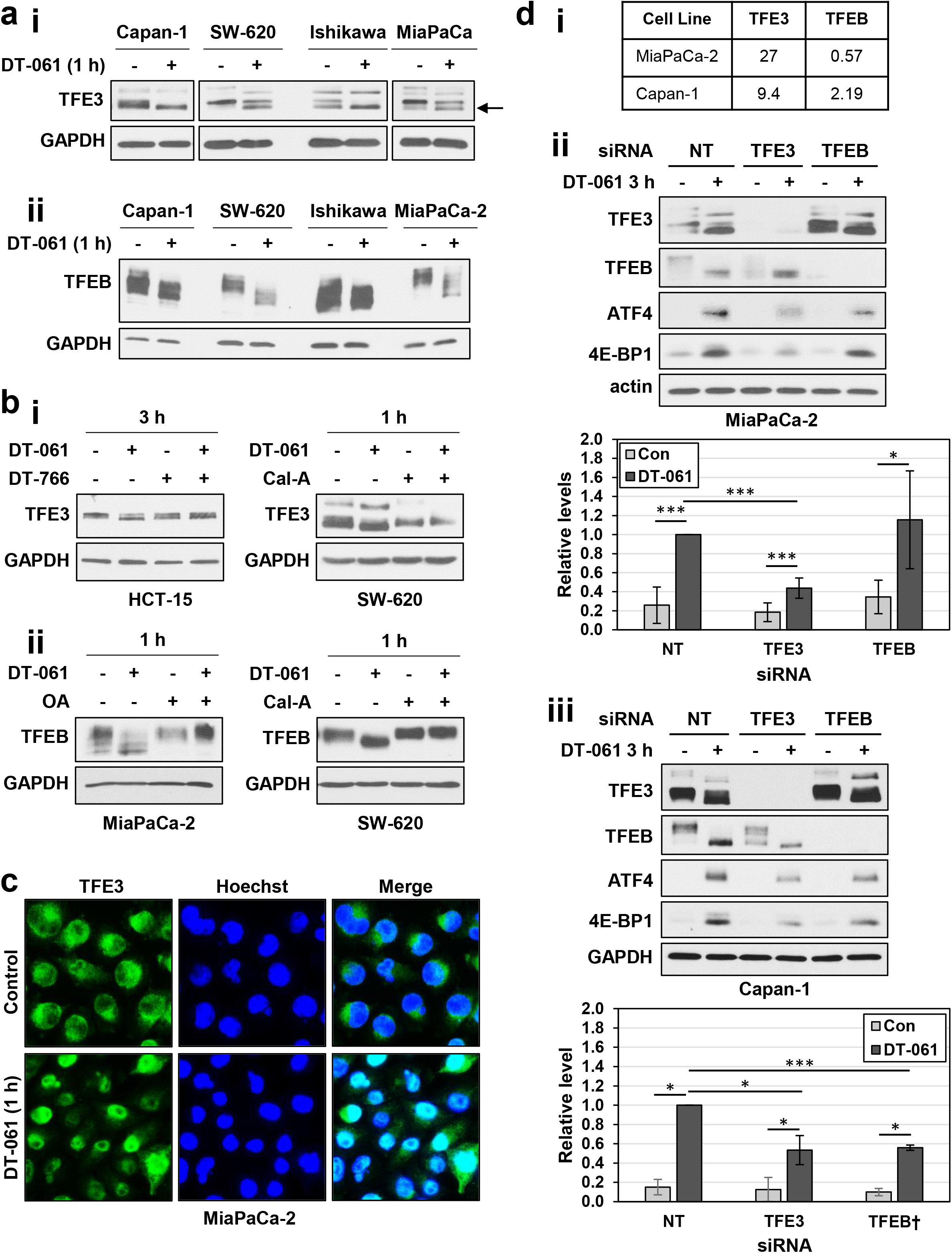
TFE3/TFEB mediate effects of SMAP/B56-PP2A on 4E-BP1 expression. ***a.i.ii.*** Cell lines were treated with vehicle or 20 µM DT-061 for 1 h before expression and electrophoretic mobility of the indicated proteins was determined by Western blotting. ***bi*** *Left panel.* Cells were treated with vehicle, 20 µM DT-061 and 80 µM DT-766 alone or in combination and analyzed for the indicated proteins by Western blotting. ***bi*** *Right panel, **bii.*** Cells were pretreated with 6 nM calyculin A (Cal-A) or 120 nM okadaic acid (OA) prior to the addition of 20 µM DT-061 for 1 h. Expression of the indicated proteins was then analyzed by Western blotting ***c***. MiaPaCa-2 cells were treated with vehicle or DT-061 for 1 h and analyzed for TFE3 subcellular localization by immunofluorescence microscopy. ***d.i.*** Relative expression of TFE3 and TFEB mRNA in the indicated cell lines (data from CCLE). ***ii,iii.*** MiaPaCa-2 or Capan-1 cells were transfected with non-targeting (NT) siRNA or siRNA targeting TFE3 or TFEB as indicated for 24 h prior to treatment with vehicle or 20 µM DT-061 for 3 h. Expression of the indicated proteins was then analyzed by Western blotting and relative signal intensity was quantified from scans of multiple exposures using Image J software (NIH). Data in *a-c* are representative of at least 3 independent experiments. Data in the graphs in *dii* and *diii* are the average ± s.e.m. of 3 independent experiments. *, p < 0.05, ***p < 0.001.

To explore the role of these factors in mediating B56-PP2A-induced 4E-BP1 upregulation, we used siRNA technology in MiaPaCa-2 and Capan-1 cells. CCLE data indicate that MiaPaCa-2 cells express approximately 50-fold more TFE3 mRNA than TFEB mRNA (Fig. 11di). In contrast, both factors are well represented in Capan-1 cells (Fig. 11di). Knockdown of TFE3 reduced the levels of SMAP-induced ATF4 upregulation in both MiaPaCa-2 and Capan-1 cells (Fig. 11dii,diii), accompanied by a significant decrease in 4E-BP1 accumulation, supporting a role for TFE3 in SMAP-induced transcriptional upregulation of 4E-BP1. Knockdown of TFEB did not significantly affect SMAP-induced upregulation of ATF4/4E-BP1 in MiaPaCa-2 cells, consistent with a predominant role for the more highly expressed TFE3 transcription factor in these cells. In contrast, TFEB knockdown reduced SMAP-induced ATF4/4E-BP1 upregulation in Capan-1 cells, indicating that TFEB is also able to mediate the effect. Together, these data point to a pathway in which B56-PP2A activation leads to dephosphorylation and nuclear accumulation of TFE3 and TFEB for transcriptional induction of ATF4 and subsequent ATF4-mediated upregulation of 4E-BP1.

## DISCUSSION

By leveraging novel small molecule activators of PP2A with selectivity for a subset of PP2A heterotrimers (29,36–39), the current study shows that B56-PP2A(s) orchestrate a translation repressive program in tumor cells involving 4E-BP1 upregulation via a transcriptional mechanism and activation of 4E-BP translation inhibitory function by hypophosphorylation (Fig. 12). Remarkably, SMAP/B56-PP2A can restore translation control in tumor cells with profound suppression of 4E-BP1 expression (e.g., many PDAC and CRC cell lines) by engaging an ATF4→4E-BP1 transcriptional axis that involves the transcription factors TFE3/TFEB (Fig. 12). Importantly, the protein that accumulates is in the active, translation repressive form as a result of B56-PP2A-mediated hypophosphorylation. SMAP/B56-PP2A inhibition of eIF4F formation and cap-dependent translation activity was confirmed by cap affinity assays, eIF4E co-IP assays, and bicistronic dual reporter cap-dependent translation activity assays.

**Figure 12.**
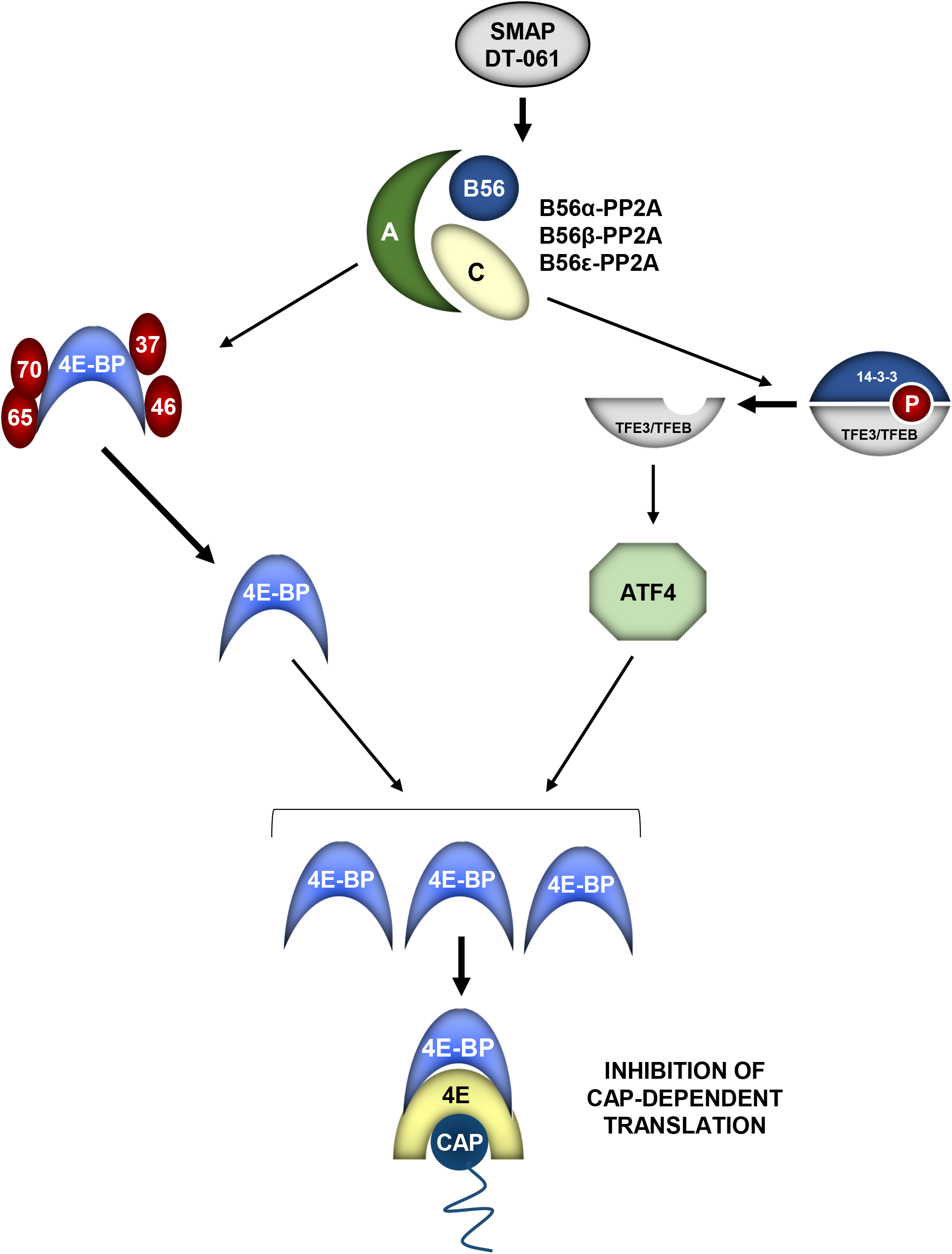
Model for the effects of SMAPs on 4E-BPs. SMAPs such as DT-061 and DT-1154 activate a subset of B56-PP2A heterotrimers (B56α-, B56β-, and/or B56ε-PP2A) to dephosphorylate 4E-BP1 and 4E-BP2 at canonical sites. SMAPs/B56-PP2A also dephosphorylate and activate the transcription factors TFEB/TFE3 to induce ATF4 transcription. ATF4 accumulation activates transcription of the 4E-BP1 gene leading to accumulation of 4E-BP1 protein that is also dephosphorylated by B56-PP2A. Active 4E-BP1 prevents formation of the eIF4F translation initiation complex and inhibits cap-dependent translation.

Since tumor cells are “addicted” to aberrant protein synthesis for reprogramming the proteome to sustain the malignant phenotype, translation control serves as an “Achilles heel” that has recently emerged as a promising therapeutic target (5,11,12). Initial efforts to target this vulnerability have focused on inhibitors of upstream 4E-BP1 kinases (18) such as mTOR, with the antitumor activity of mTOR inhibitors largely reflecting their ability to abrogate 4E-BP1 phosphorylation. However, numerous studies have shown that resistance to mTOR inhibitors arises through 4E-BP1 phosphorylation by the ERK pathway (18,20,65). Similarly, ERK-independent 4E-BP1 phosphorylation underlies resistance to anti-BRAF and anti-MEK therapies in CRC and other cancers (21). Thus, strategies for suppression of aberrant cap-dependent translation that do not rely on inhibition of upstream 4E-BP kinases offer therapeutic promise. Intense efforts are currently focused on directly targeting the eIF4F complex by suppressing eIF4E with antisense oligonucleotides (ISIS-EIF4ERx) (66), disrupting eIF4F complex formation with inhibitors of the eIF4E/eIF4G interaction (e.g., 4EGI-1, 4E1RCat, 4E2RCat) (67), impeding eIF4E–cap binding with 4Ei-1 (68), and inhibiting the eIF4A helicase with silvestrol or rocaglamide A (5,69–71). Our discovery that SMAPs block eIF4F assembly and function through PP2A-dependent activation of 4E-BP1 points to an additional mechanism for restoration of translation control in cancer. Importantly from a therapeutic perspective, our data further demonstrate that SMAP/B56-PP2A is able to activate 4E-BPs in the context of common mechanisms of PP2A inhibition in cancer cells [e.g., overexpression of endogenous PP2A inhibitors SET (Capan-1) and CIP2A (MiaPaCa-2) or R183 mutation of the A scaffolding subunit (HCT-116) (72–74)]. Thus, the findings of the current study highlight the potential of PP2A-based strategies to restore the function of 4E-BPs and repress aberrant cap-dependent translation for therapeutic benefit.

Our study provides the first evidence that B56-PP2A heterotrimer(s) can activate 4E-BP1 and 4E-BP2 by hypophosphorylation. B56-PP2A reduces phosphorylation at Thr37, Thr46, Thr70 and Ser65 and thus targets all major 4E-BP sites. While previous studies had linked PP2A to regulation of 4E-BP phosphorylation (47), the regulatory subunit(s) responsible for 4E-BP targeting had not been defined. Our knockdown studies support a role for B56α, a major regulatory subunit targeted by SMAPs (37), although our data suggest that B56β and/or B56ε may also be able to mediate the effect. Our analysis indicates that SMAP/B56-PP2A can hypophosphorylate 4E-BPs in the presence of active upstream negative regulators such as mTOR, AKT, and ERK, supporting a role of B56-PP2A(s) as 4E-BP phosphatases. This conclusion is consistent with the demonstration that FLAG-tagged PP2A-C can dephosphorylate GST-4E-BP1 *in vitro* (24), although the ability of B56-PP2A(s) to interact with 4E-BPs remains to be confirmed.

Based on the known function of SMAPs as activators of PP2A, a major goal of this study was to test DT-061 for the ability to dephosphorylate 4E-BPs and thus reestablish homeostatic translation control in cancer via activation of hyperphosphorylated/inactivated 4E-BPs that are commonly overexpressed in tumor cells. The finding that SMAPs can also restore and upregulate the expression of 4E-BP1 via a transcriptional mechanism was unexpected. Our data show that B56-PP2A can rapidly induce the expression of the transcription factor ATF4 at the mRNA and protein levels, and that ATF4 plays a requisite role in B56-PP2A-mediated transcriptional upregulation of 4E-BP1. Consistent with previous studies showing that ATF4 can induce the expression of 4E-BP1 (58,60), but not 4E-BP2 or 4E-BP3, SMAPs only affected the expression of 4E-BP1.

SMAP-induced upregulation of ATF4 was blocked by the transcription inhibitor ActD, indicating that B56-PP2A enhances transcription of the ATF4 gene. ATF4 has recently been identified as a direct target of the transcription factors TFE3/TFEB, which recognize a CLEAR motif (GT<u>CACGTG</u>AC) in the ATF4 promoter located 875 bp upstream of the transcription start site (61). Notably, SMAP/B56-PP2A also promotes hypophosphorylation of TFE3/TFEB. Dephosphorylation of these factors is required for their translocation to the nucleus for activation of transcription, with calcineurin and B55α-PP2A recently identified as TFE3/TFEB phosphatases (61–63). Use of SMAPs provides the first evidence that B56-PP2A may also dephosphorylate these factors, allowing them to translocate to the nucleus and activate transcription. Our demonstration that knockdown of TFE3/TFEB reduces the ability of SMAP/PP2A to upregulate 4E-BP1 in multiple cell lines provides the first evidence for regulation of 4E-BP1 expression by these factors, while suggesting a model in which B56/PP2A activates TFE3/TFEB by dephosphorylation, and TFE3/TFEB in turn induce the expression of ATF4 for transcriptional upregulation of 4E-BP1 (Fig. 12). Ongoing studies are testing this model.

4E-BP translation inhibitors function as tumor suppressors and tumorigenesis is critically dependent on inhibition of 4E-BP activity. Many tumors express abundant levels of inactive hyperphosphorylated 4E-BP1, which is available for reactivation and tumor suppression. Other tumors potently suppress 4E-BP expression or overexpress eIF4E to increase the eIF4E:4E-BP ratio and overwhelm the inhibitory capacity of 4E-BPs. By upregulating and hypophosphorylating 4E-BP1, B56-PP2A activation is a promising approach for restoration of translation control in tumors harboring different mechanisms of 4E-BP inactivation.

## Materials and Methods

### Cell Culture

Colon cancer cell lines: HCT-15 (ATCC CCL-225), RKO (ATCC CRL-2577), LoVo (ATCC CCL-229), SW-620 (ATCC CCL-227), SW-480 (ATCC CCL-228), FET (75) and HCT-116 (ATCC CCL-247) were maintained in RPMI 1640, 10% fetal bovine serum, 2 mM L-glutamine, and penicillin/streptomycin. Pancreatic cancer cell lines: Capan-1 (ATCC HTB-79), Capan-2 (ATCC HTB-80), Panc-1 (ATCC CRL-1469), BxPC-3 (ATCC CRL-1687), MiaPaCa-2 (ATCC CRM-CRL-1420), Panc89 (RRID:CVCL_4056; a gift from Dr. M.A. Hollingsworth, UNMC), and S2-013 (RRID:CVCL_B280; from M.A. Hollingsworth, UNMC) were maintained in RPMI 1640, 10% fetal bovine serum, 2 mM L-glutamine, and penicillin/streptomycin. The human endometrial cancer cell line Ishikawa (RRID:CVCL_2529) was a gift from Dr. Tim Hui-Ming Huang (Ohio State University). Ishikawa cells were maintained in DMEM, 10% fetal bovine serum, 1% non-essential amino acids, 5 μg/ml insulin and penicillin/streptomycin. The human endometrial cancer cell line SNG-M (RRID:CVCL_1707, from the Japanese Collection of Research Bioresources Cell Bank) was maintained in Ham’s F-12, 10% fetal bovine serum and penicillin/streptomycin. H358 non-small cell lung cancer cells (ATCC CRL-5807) were maintained in RPMI 1640, 10% fetal bovine serum, and 0.5% penicillin/streptomycin. IEC-18 cells (ATCC CRL-1589) were maintained in Dulbecco’s modified Eagle’s medium (Invitrogen) with 4 mM glutamine, 5 μg/ml insulin, and 5% fetal bovine serum. All cell lines were maintained in 5% CO_2_ at 37°C.

### Drug Treatments

Cells were treated for the indicated times with 20 μM DT-061 (or as indicated), 20 μM DT-1154, 10 nM INK128 (Cayman Chemical Company), 20 μM DT-1310, 80 μM DT-766, 6 nM Calyculin A (Sigma), 120 nM okadaic acid (Millipore Sigma), or 1 μg/ml actinomycin D (Sigma). All drugs were resuspended in DMSO with the exception of okadaic acid, which was resuspended in 50% DMSO/50% ethanol. Equal volumes of vehicle were added to all control plates, and final solvent concentrations were ≤ 0.2%.

### Western Blotting

Cells were lysed in 1% SDS, 10 mM Tris-HCl, pH 7.4, and equal amounts of protein were subjected to SDS-PAGE and transferred to nitrocellulose membrane as we have described previously (76). Equal loading and even transfer was confirmed by staining the membranes with 0.1% fast green (Sigma-Aldrich). Membranes were blocked for 30 minutes in Tris-buffered Saline, 0.1% Tween-20 (TBS-T) containing 5% non-fat dried milk or 5% BSA. Membranes were then incubated with primary antibody overnight at 4 °C, washed and incubated with horseradish peroxidase-conjugated secondary antibody for 2 h at room temperature. Following extensive washing, signal was detected using SuperSignal West Pico Chemiluminescence Substrate (ThermoScientific) and CL-XPosure Film (ThermoScientific). Relative signal intensity of bands was quantified from scans of multiple exposures using Image J software (NIH).

### Antibodies

Primary antibodies were obtained from Cell Signaling Technology (CST), Santa Cruz Biotechnology (sc), Abnova, Sigma-Aldrich, AbClonal, and ProteinTech as indicated. Dilutions used in Western blotting were as follows: 4E-BP1 (CST-9644, 1:20,000), 4E-BP2 (CST-2845, 1:2,000 to 1:4,000), 4E-BP3 (Abnova, H00008637-M05, 1:500), phospho-64/65 4E-BP1 (CST-9451, 1:500 and sc-18091-R, 1:500), non-phospho-45/46 4E-BP1 (CST-4923, 1:2,000 to 1:4,000), GAPDH (CST-5174, 1:60,000), Actin (Sigma A2066, 1:20,000 or AbClonal AC026, 1:10,000-20,000), eIF4E (sc-271480 or sc-9976, 1:2,000 to 1:4,000), phospho-eIF4E Ser209 (CST-9741, 1:1,000), eIF4G (sc-133155, 1:1000 or CST-2469, 1:100,000), total AKT (CST-9272, 1:3,000), phospho-AKT S473 (CST-4060, 1:20,000), phospho-AKT T308 (CST-2965, 1:1000 to 1:8,000), pULK-1 (CST-14202, 1:1,000), total p70S6K (sc-8418, 1:2,000), phospho-p70S6K (AbClonal AP0564, 1:2,000), total ERK (CST-9102, 1:2,000), phospho-ERK (CST-9106, 1:2,000), B56α (sc-136045, 1:200), B55α (CST-2290, 1:2000), ATF4 (Proteintech, 10835-1-AP, 1:2,000), TFE3 (CST-14779, 1:2,000), and TFEB (CST-4240, 1:1,000). All primary antibodies were suspended in Tris-buffered Saline with 5% BSA except for actin and eIF4E, which were diluted in TBS-T with 5% non-fat dried milk. Goat anti-rabbit-HRP antibody (Millipore, AP132P) was used at 1:1,000 to 1:2,000, goat anti-rabbit-HRP antibody (Jackson Immunoresearch Laboratories, 111-035-144) was used at 1:10,000 to 1:50,000, and goat anti-mouse-HRP antibody (Bio-Rad, 170-6516) was used at 1:3,000. All secondary antibodies were diluted in Tris-buffered Saline with 5% milk.

### RNA Interference

For ATF4, TFEB, and TFE3 knockdown, cells were transfected with 100 pmol siRNA using RNAiMAX transfection reagent (Invitrogen) and cells were analyzed after 24 h. PPP2R5A/B56α and PPP2R2A/B55α were knocked down using 100 pmol of siRNA for a total of 48 or 72 hours before drug treatment and analysis. siRNAs from Ambion Life Technologies were as follows: ATF4 siRNA#1, GUGCUGUAGCUGUGUGUUC; ATF4 siRNA#2, CCUGGAAACCAUGCCAGAU; TFEB siRNA, AGACGAAGGUUCAACAUCA; TFE3 siRNA, GGCGAUUCAACAUUAACGA; B55α (Rat and Human specific) Silencer Select Custom siRNA ID # s554710, UAAAACUCCUGUCUGUAAUtt. siRNAs from Dharmacon were as follows: Human PPP2R5A ON-TARGET plus SMARTpool, L-009352-00-0005; Rat PPP2R5A ON-TARGET plus SMARTpool, L-083443-02-0005. Controls were transfected with equivalent levels of ON-TARGET plus non-targeting siRNA (Dharmacon, D-001810-01-05).

### Cap Affinity Assay

Cap affinity chromatography was performed as previously described (77). SMAP-or vehicle-treated cells were harvested in cap affinity lysis buffer (50 mM Tris, pH 7.4, 100 mM NaCl, 1 mM EDTA, and 1% Triton X-100, supplemented with protease and phosphatase inhibitor cocktails), and particulate material was removed by centrifugation. Cell lysates (500 mg of protein) were incubated with 50 μl of 7-methyl-GTP-Agarose slurry (Jena Biosciences, AC-155S) for 16 h at 4 °C. Beads were washed four times with cap affinity lysis buffer, and cap-bound protein was eluted with 25-50 μl of 2x SDS sample buffer and subjected to Western blot analysis.

### Immunoprecipitation

Immunoprecipitation was performed as we have described (26). Briefly, cells were lysed in IP lysis buffer (50 mM Tris-HCl, pH 7.5, 150 mM NaCl,1 mM EDTA, 1% IGEPAL, 10% glycerol, Protease Inhibitor Cocktail and Phosphatase Inhibitor Cocktails I and II (Sigma)). Extracts were incubated with 1-2 μg of anti-eIF4E (sc-271480) antibody for 16 h at 4°C before protein G-PLUS-agarose slurry (Santa Cruz sc-2002) was added for 2 h at 4°C. Beads were washed 4 times in IP lysis buffer, resuspended in 25-50 μl 2x SDS-PAGE sample buffer and analyzed by Western blotting.

### Bicistronic cap-dependent translation activity assay

MiaPaCa2 cells were transfected with the bicistronic pcDNA3-RLUC-POLIRES-FLUC plasmid (Addgene 45642) (55) using Lipofectamine 3000 (ThermoFisher) and selected with 0.8 mg/ml G418. For analysis of effects on cap-dependent translation, pooled stable transfectants were plated at 8.33-16.66 x 10^4^ cells per well in six well plates in the absence of G418 48 h prior to treatment with 20 µM DT-061, 500 nM INK128 or vehicle. After 6 h of treatment, firefly and Renilla luciferase activities were quantified using the Dual Luciferase Reporter Assay System (Promega Corp., Madison, WI) and a Berthold Lumat LB 9501 luminometer. Relative cap-dependent translation was derived by normalizing Renilla luciferase activity to firefly luciferase activity for each sample.

### Reverse Transcription Quantitative PCR (RT-qPCR)

Total cellular RNA was isolated using Trizol reagent (ThermoFisher) and RT-qPCR was performed using the 1-Step Brilliant II SYBR Green quantitative RT-PCR master mix kit (Agilent Technologies) on a Bio-Rad C1000 Thermal Cycler CFX96 Real-Time System. Relative mRNA concentrations were determined using CFX Manager Software, v3.1 (BioRad) from standard curves, and mRNA levels were normalized to 18 S rRNA levels for each sample. Primers used for RT-qPCR and qPCR were as follows: Human 4E-BP1, GGAGTGTCGGAACTCACCTG and ACACGATGGCTGGTGCTTTA; Rat 4E-BP1, ACACAGAGGAGTCTGTCGGA and AAGGTAAGGTGGGTGTGCTC; Human ATF4, TTCTCCAGCGACAAGGCTAAGG and CTCCAACATCCAATCTGTCCCG; Human and Rat 18S rRNA, CATTGGAGGGCAAGTCTGGTG and CTCCCAAGATCCAACTACGAG; Human GAPDH, TGAAGGTCGGAGTCAACGGA and CCATTGATGACAAGCTTCCCG.

### Nascent RNA Capture

Labeling and capture of nascent RNA were performed using the Click-iT Nascent RNA Capture Kit (Life Technologies) as instructed by the manufacturer. In brief, 5-ethynyl uridine (EU) was added to the culture medium (0.5 mM) for the final 1 h of drug/vehicle treatment and cellular RNA was purified using Trizol reagent (ThermoFisher). Newly synthesized, EU-labeled RNA was biotinylated using Click chemistry and isolated using Streptavidin magnetic beads. cDNA was generated from bead-bound EU-labeled RNA using the iScript cDNA Synthesis Kit (Bio-Rad Laboratories) and analyzed by qPCR using iTaq Universal SYBR Green Supermix (Bio-Rad Laboratories). Normalization to nascent GAPDH mRNA and calculation of relative levels used the ΔΔC_t_ method.

### Proteomic analysis of 4E-BP1 expression

PDAC tissues and corresponding tumor-adjacent tissue were prepared for LC/MS-MS and analyzed on a Dionex Nano Ultimate 3000 coupled with an Orbitrap Fusion Lumos as described (78). Bioinformatic analysis was as described (78).

### Phosphoproteomic analysis

Samples were lysed in 2% SDS, subjected to a2-step Lys-C/trypsin proteolytic cleavage, and phophopeptides were enriched using titanium dioxide spin tips (Thermo Fisher, IL). LC-MS/MS analysis was performed on an Orbitrap ProVelos Elite MS system (Thermo Fisher Scientific) equipped with a nanoAcquity ultra-high pressure liquid chromatography system (Waters, MA). Peptide clustering and phosphopeptide quantification was performed as described previously (79,80).

### Immunofluorescence analysis

Cells were plated onto pre-sterilized glass coverslips at a density of 1x10^5^ cells and cultured overnight. Cells were then treated with vehicle (DMSO) or 20 μM DT-061 for 1 h, followed by fixation in 4% formaldehyde/PBS for 15 minutes at room temperature. Coverslips were then processed for immunofluorescence microscopy as we have described previously, using 0.2% Triton X-100 for permeabilization. TFE3 (CST 14779) primary antibody was used at 1:100 and Alexa Fluor 488-conjugated donkey anti-rabbit secondary antibody was used at 1:1,000. Nuclei were stained with Hoechst 33258. Images were obtained using a Zeiss Axio Observer.Z1 microscope with a Zeiss Axiocam 208 Color digital camera.

### Analysis of Publicly Available Data Sets

Analysis of 4E-BP1 mRNA and protein expression in normal and tumor tissue using The Cancer Genome Atlas (TCGA) and Clinical Proteomic Tumor Analysis Consortium (CPTAC) data, respectively, was performed using the University of Alabama at Birmingham UALCAN interactive web portal (http://ualcan.path.uab.edu/analysis.html) (81).

### Statistical analysis and other software

Numerical data are presented as means ± standard error of the mean (s.e.m.) or standard deviation (s.d.) as appropriate. For presentation, contrast and brightness of scanned images were adjusted using GMU Image Manipulation Program (GIMP), Adobe Photoshop, or Microsoft PowerPoint Software. All adjustments to contrast and brightness were made equally across the entire blot and no individual lanes were treated differently than the rest of the blot. Places in images where lanes have been rearranged for clarity are indicated by dashed lines. Graphs were generated using Microsoft Excel Software and figures were assembled and annotated in Microsoft PowerPoint and Photoshop Software. Other software used is listed above.

## DATA AVAILABILITY

All data described in the manuscript are contained within the manuscript or in Supporting Information.

## SUPPORTING INFORMATION

This article contains supporting information.

## ACKNOWLEDGMENTS

Data used in this publication were generated by the TCGA Research Network (https://www.cancer.gov/tcga) and the Clinical Proteomic Tumor Analysis Consortium (NCI/NIH).

## AUTHOR CONTRIBUTIONS

**Michelle Lum:** Investigation, Methodology, Data Curation, Formal Analysis, Writing-Original draft preparation, Reviewing and Editing; **Kayla Jonas-Breckenridge:** Investigation, Data Curation, Formal Analysis, Writing-Original draft preparation, Reviewing and Editing; **Adrian Black**: Conceptualization, Investigation, Methodology, Data Curation, Formal Analysis, Writing-Original draft preparation; Reviewing and Editing, Supervision, Visualization; **Nicholas T. Woods:** Investigation, Formal Analysis; **Caitlin O’Connor:** Data Curation, Formal Analysis; **Rita Avelar:** Investigation; **AnaLisa DiFeo:** Conceptualization, Investigation, Formal Analysis**; Goutham Narla:** Conceptualization, Investigation, Formal Analysis, Reviewing and Editing, Funding Acquisition; **Jennifer Black**: Conceptualization, Investigation, Methodology, Formal Analysis, Writing-Original draft preparation, Supervision, Visualization, Writing, Reviewing and Editing, Funding Acquisition.

## SUPPLEMENTAL FIGURE LEGENDS

**Figure S1. SMAPs do not induce upregulation of 4E-BP3.** The indicated cells were treated with vehicle or 20 µM DT-061 for 3 h and expression of actin and 4E-BP3 was analyzed by Western blotting. Rap: Extracts of MiaPaCa-2 cells treated with rapamycin for 24 h to upregulate 4E-BP3 (50) used as a positive control for detection of 4E-BP3.

**Figure S2. Chemical structure of DT-061, DT-1154, DT-1310, and DT-766.**

**Figure S3. ATF4 Chip-seq data in K562 and HepG2 cells from the ENCODE database.** The number in parentheses above the peak represents its distance relative to the TSS.

**Figure S4. B55α-PP2A does not mediate the effects of SMAPs on TFE3 and TFEB.** Cell lines were transfected with non-targeting (NT) or B55α targeting siRNA 72 h prior to treatment with vehicle or 20 µM DT-061 for 1 h, and expression and electrophoretic mobility of the indicated proteins were determined by Western blotting.

