## Supplemental Figures S1-S4 for "Small Molecule Activators of B56-PP2A Restore 4E-BP Expression and Function to Suppress Cap-dependent Translation in Cancer Cells"

### Supplemental Figure S1

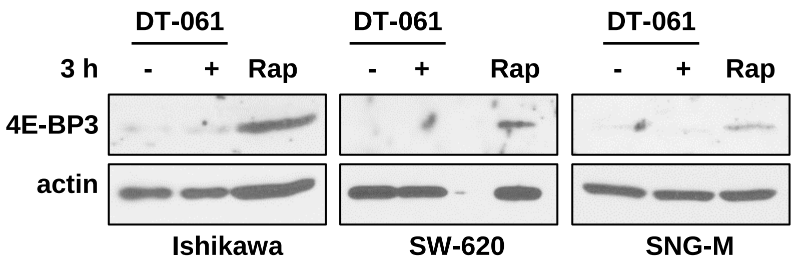


**Figure S1. SMAPs do not induce upregulation of 4E-BP3.** The indicated cells were treated with vehicle or 20 µM DT-061 for 3 h and expression of actin and 4E-BP3 was analyzed by Western blotting. Rap: Extracts of MiaPaCa-2 cells treated with rapamycin for 24 h to upregulate 4E-BP3 (50) used as a positive control for detection of 4E-BP3.

#### Supplemental Figure S2

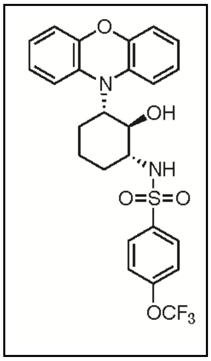

DT-061

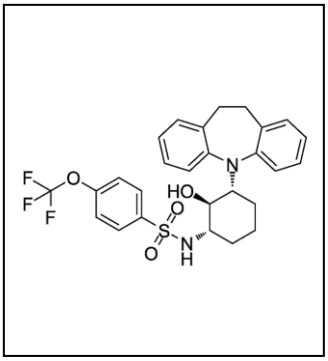

DT-1154

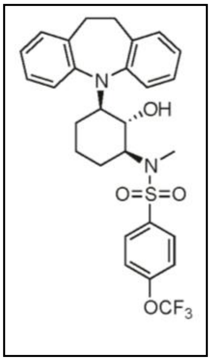

DT-1310

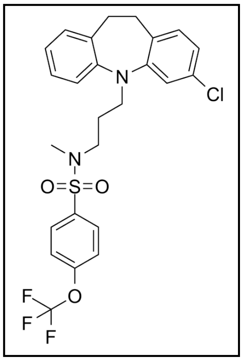

DT-766

**Figure S2. Chemical structure of DT-061, DT-1154, DT-1310, and DT-766.**

### Supplemental Figure S3

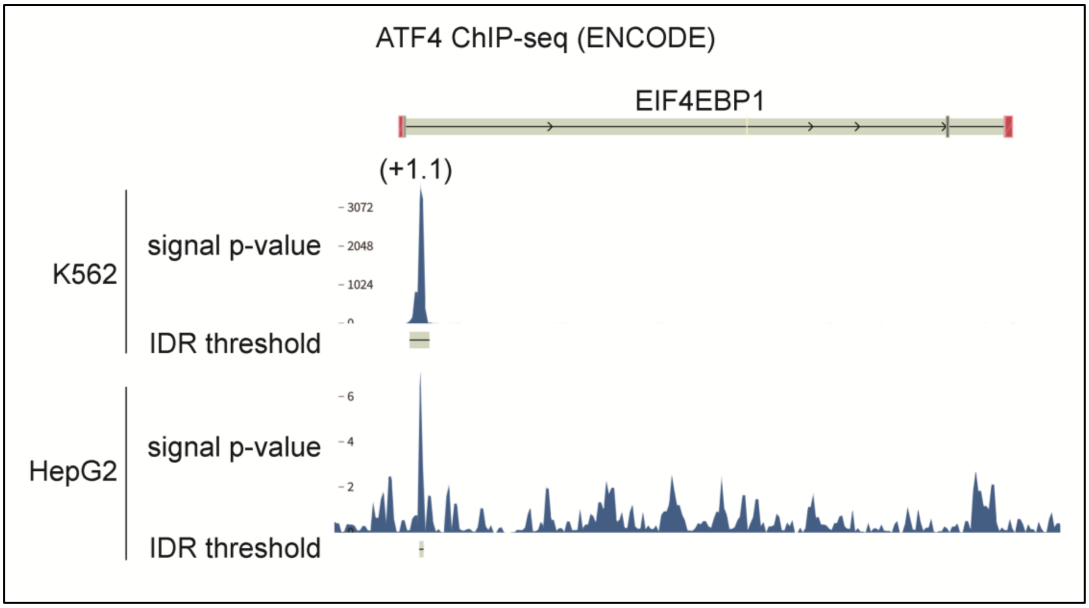


**Figure S3.** ATF4 Chip-seq data in K562 and HepG2 cells from the ENCODE database. The number in parentheses above the peak represents its distance relative to the TSS.

### Supplemental Figure S4

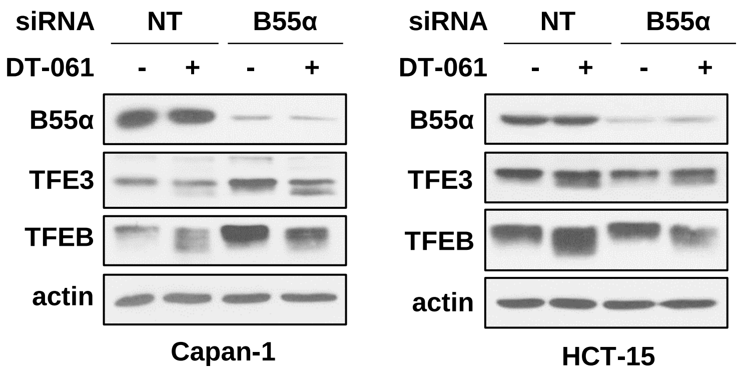


**Figure S4. B55α-PP2A does not mediate the effects of SMAPs on TFE3 and TFEB.** Cell lines were transfected with non-targeting (NT) or B55α targeting siRNA 72 h prior to treatment with vehicle or 20 µM DT-061 for 1 h, and expression and electrophoretic mobility of the indicated proteins were determined by Western blotting.
